# Suppression of T cell function by phosphoethanolamine, a metabolite enriched in tumor interstitial fluid

**DOI:** 10.1101/2024.06.10.598334

**Authors:** Yupeng Wang, Drew Wilfahrt, Chufan Cai, Konstantinos Lontos, Benjamin Cameron, Bingxian Xie, Ronal M. Peralta, Roya AminiTabrizi, Hardik Shah, Dayana B. Rivadeneira, Alexander Muir, Greg M. Delgoffe

## Abstract

Nutrient stress represents a significant barrier for antitumor immunity, and tumor interstitial fluid (TIF) often contains metabolites that hinder immune function. However, it is difficult to isolate the effects of tumor nutrient stress from other suppressive factors. Thus, we employed a chemically-defined cell culture medium based on the metabolomic profile of TIF: Tumor Interstitial Fluid Medium (TIFM). Culture of CD8^+^ T cells in TIFM limited cell expansion and impaired CD8^+^ T cell effector functions upon restimulation, suggesting tumor nutrient stress alone is sufficient to drive T cell dysfunction. We identified phosphoethanolamine (pEtn), a phospholipid intermediate, as a driver of T cell dysfunction. pEtn dampened TCR signaling by depleting T cells of diacylglycerol required for TCR signal transduction. Reduction of pEtn accumulation in tumors improved intratumoral T cell function and tumor control, suggesting pEtn accumulation plays a dominant role in TME immunosuppression.

## Introduction

T cells are a critical component of antitumor immunity: higher infiltration of T cells in solid tumors is associated with more effective tumor control^1,2^, and checkpoint blockade therapies using antibodies targeting Programmed Death 1 (PD-1) and other inhibitory receptors have achieved durable responses in the clinic^3^. Still, many patients fail to respond to these therapies. Often, non-responders have dysfunctional tumor-infiltrating T cells^4–6^. Ultimately, intratumoral T cells have decreased effector functions and are unable to mount a proliferative burst needed to control tumors. Multiple non-redundant mechanisms drive immune dysfunction in the TME^7^. Chronic exposure to tumor antigen, especially in the context of TME metabolic stress, has been shown to induce T cell exhaustion, a hypofunctional T cell state^8,9^. In addition, T cells also receive inhibitory signals in the TME from suppressive cytokines and ligands to inhibitory receptors that limit immune functionality^10^. Lastly, tumor-derived metabolic stresses including hypoxia, nutrient depletion, and exposure to accumulated metabolic waste can render T cells metabolically impaired and dysfunctional^11,12^. Understanding how each of these TME components limit immune control of tumors and by what mechanisms can provide new avenues to intervene and improve immunotherapeutic outcomes.

It is now clear that T cell acute effector functions, proliferation, and differentiation can be profoundly influenced by changes in metabolism^13,14^. While naive T cells are metabolically quiescent, upon activation T cells initiate many metabolic processes including lipid synthesis to support proliferation, differentiation, and effector function^15^. These enhanced metabolic activities of effector T cells require sufficient supply of extracellular nutrients. Tumor cells consume local nutrients and release excessive metabolic byproducts^16^. Additionally, metabolic activity in stromal and immune cells prevalent in tumors can drive both nutrient deprivation and accumulation of suppressive metabolic byproducts^17–20^. The suppressive metabolic activity in cancerous and tumor-associated cells is further exacerbated by the insufficient metabolic exchange between blood and the interstitial fluid due to dysfunctional vasculature in the tumor^21^. Therefore, the imbalance of helpful nutrients and harmful waste could skew the metabolism of T cells and contribute to T cell dysfunction in the TME. Restoration of relative balance in the nutrients landscape in the TME can enhance anti-tumor immunity^22,23^. However, the complex composition of the metabolites and other suppressive factors in the TME make it difficult to isolate the effects of tumor nutrient stress on immune cells and identify all the key TME metabolic drivers of T cell dysfunction.

Recently, we developed quantitative metabolomics to measure the concentration of over 118 metabolites in the tumor interstitial fluid of murine pancreatic and lung tumors^24^. These studies provided novel insight into the nutritional composition of the TME in solid tumors. Further, we developed a cell culture medium formulation termed Tumor Interstitial Fluid Medium (TIFM) containing TME levels of 115 major nutrients, allowing us to model tumor nutrient stress in vitro^25^. In this study, we used TIFM to study the effects of environmental nutrient availability on T cell biology and function, providing a platform to study how T cells react to TME nutrient stress without interference from other confounding immunosuppressive signals in the TME. We show that CD8^+^ T cells develop dysfunctional features during culture in TIFM: increased expression of PD-1, decreased cytokine production, and impaired tumor cell killing. Thus, TME nutrient stress is sufficient to drive T cell dysfunction. By manipulating metabolite levels in TIFM, we identified key metabolites that are depleted in the TME that are essential for T cell proliferation, and a novel oncometabolite enriched in multiple cancers that can drive T cell dysfunction.

## Results

### Culture in TIFM induces robust CD8^+^ T cell dysfunction

Based on previously reported measurements of the metabolome of tumor interstitial fluid, we generated a chemically defined media formulation containing 115 metabolites at TME levels, termed Tumor Interstitial Fluid Medium (TIFM) (Table S1)^25^ which we supplemented with 10% dialyzed FBS to provide growth factors and lipids required for T cell growth and function. TIFM contains salts and sodium bicarbonate at RPMI-1640 (termed RPMI herein) levels, making TIFM and RPMI identical except in nutritional content, allowing us to isolate the effects of tumor-derived nutrient stress on T cell physiology. Thus, we used TIFM to determine how T cells respond to the metabolite imbalances experienced by the T cell during tumor infiltration.

Given the importance of the nutrient environment on T cell function and differentiation^13^, we first assessed how CD8^+^ T cells responded when cultured for an extended period in TIFM and RPMI after activation. OT-I T cells were activated for 24 hours with cognate peptide in complete RPMI-based growth media (RPMI supplemented with 10% fetal bovine serum, antibiotic/antimycotics, and sodium pyruvate). After activation, cells were cultured for 24 additional hours in RPMI supplemented with murine IL-2 and then transferred into TIFM for 5 days prior to day 7 analysis (Figure 1A). An additional group was kept in control RPMI for all 7 days of cell culture, which serves as a standard T cell culture control. Upon exposure to TIFM, T cells stopped robustly proliferating (Figure 1B). Further, CD8^+^ T cells cultured in TIFM for 5 days had significantly elevated expression of PD-1 compared to RPMI cultured cells. (Figure 1C). We then sought to determine whether TIFM-cultured cells had impaired effector functions compared to RPMI. T cells on day 7 were restimulated with anti-CD3 and anti-CD28 in RPMI (regardless of their prior culture conditions) and cytokine production was measured by flow cytometry after 6 hours of restimulation (Fig 1A). T cells cultured in TIFM for 5 days prior to restimulation had fewer polyfunctional cells that express both IFNγ and TNF after restimulation than RPMI controls (Figure 1D). Additionally, T cells exposed to TIFM had a reduction in Granzyme B protein levels compared to RPMI-cultured controls, while IL-2 levels were not significantly different between groups (Figures 1E, 1F). Given these observations, we hypothesized that T cells cultured in TIFM and RPMI would have different capacities for tumor cell killing. To test this, OT-I T cells were activated, and then cultured as before in physiologic nutrient conditions with TIFM for 5 days. On day 7 of culture cells were placed into *in vitro* killing assays with B16 melanoma cells expressing ovalbumin (B16-OVA) and monitored for tumor cell killing over time using impedance with an xCelligence RTCA DP device. RPMI-cultured CD8^+^ T cells were able to achieve nearly 100% cytolysis after 30 hours in the co-culture assay (Figure 1G). Strikingly, T cells cultured in TIFM for 5 days prior to the assay killed tumor cells far more slowly and failed to kill all tumor cells (Figure 1G), indicating that prolonged exposure to tumor nutrient stress impaired CD8^+^ T cell killing of target tumor cells. These data show that the tumor nutrient environment induces a unique cell surface and cytokine profile with elevated PD-1 expression and reduced IFNγ and TNF cytokine expression, and impaired tumor cell killing capacity.

**Figure 1:**
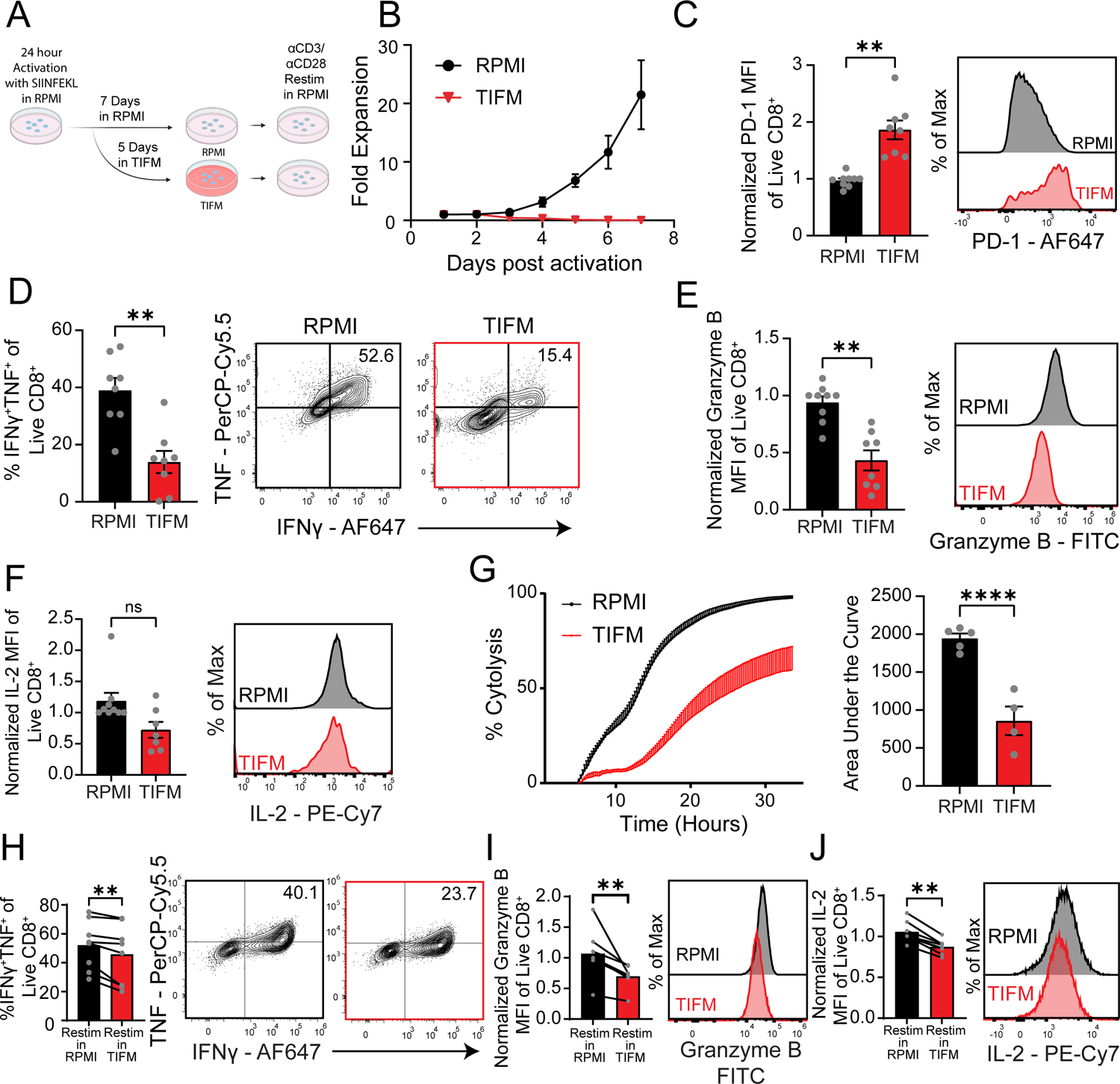
Culture in TIFM induces CD8^+^ T cell dysfunction **A)** Splenocytes from OT-I mice were activated with peptide for 24 hours in complete RPMI. Cells were cultured for an additional 24 hours in complete RPMI supplemented with IL-2, and then transferred to Tumor Interstitial Fluid Like Medium (TIFM) with IL-2 for 5 days. Control cells were cultured in RPMI throughout. After 7 days, all cells were restimulated in RPMI with 3µg plate-bound αCD3 and 1µg soluble αCD28 antibody for 6 hours. **B)** Fold expansion from cell culture conditions described in A). Plot shows fold expansion ± SEM. (n=4/group) **C)** Expression of PD-1 from Day 7 OT-I CD8^+^ T cells from indicated culture conditions and quantification of normalized PD-1 MFI (mean fluorescence intensity) ± SEM from four independent experiments is shown on the left (n=8-9/group). **D)** Day 7 OT-I CD8^+^ T cells were analyzed for IFNγ and TNF co-expression after 6 hours αCD3/αCD28 restimulation. Data quantified on the left show %IFNγ^+^TNF^+^ events among Live CD8^+^ T cells ± SEM from four independent experiments (n=8/group). **E-F)** Day 7 OT-I T cells were analyzed for expression of **E)** Granzyme B or **F)** IL-2 after 6 hours of αCD3/αCD28 restimulation. Data quantified on the left show normalized MFI ± SEM from four independent experiments (n=8-9/group). **G)** Day 7 OT-I CD8^+^ T cells were co-cultured with B16-OVA cells at a 1:1 effector:target ratio in an Agilent xCELLigence RTCA DP system to monitor target cell killing. XY plot on the left shows % Cytolysis ± SEM, and Area under the curve values are quantified on the right from two independent experiments (n=4-5/group). Statistical analysis for all indicated comparisons in **C-G)** was determined using a Wilcoxon matched-pairs signed rank test. **H-J)** OT-I Splenocytes were cultured in RPMI for 7 days as described in **A)** and then restimulated with αCD3/αCD28 in RPMI or TIFM for 6 hours. After restimulation, cells were analyzed for **H)** IFNγ and TNF co-expression **I)** Granzyme B expression and **J)** IL-2 expression. MFI values from multiple experiments were normalized and are quantified on the left (n=8/group from three independent experiments). For **H-J)**, statistical significance was determined using a Wilcoxon matched-pairs signed rank test. For indicated comparisons, * = p<0.05, ** = p <0.01, *** = p<0.001, and **** = p<0.0001.

Importantly, the effects seen in these experiments are dependent upon exposure time to each nutrient environment. Cells cultured in TIFM for 1 or 3 days, respectively, had less pronounced phenotypes than those cultured in tumor nutrient stress for 5 days (Figure S1A-F). Cells cultured in TIFM for 3 days still had a significant increase in PD-1 expression compared to RPMI-cultured cells, but no statistically significant differences were seen for IFNγ and TNF co-expression or IL-2 expression (Figure S1C-S1D). Granzyme B expression was still significantly reduced in cells cultured in TIFM for at least 1 day (Figure S1E), consistent with 5-day culture results (Figure 1E). Cell killing was significantly reduced in the cells cultured in TIFM for 3 days, but 1 day in TIFM did not impair tumor cell killing capacity (Figure S1F).

For our initial experiments (Figure 1A-G), T cells were restimulated in RPMI to control for potential nutrient-specific effects on TCR signaling previously described^26^. Still, we wanted to test whether short-term exposure to tumor nutrient stress during restimulation had an effect on T re-activation. To test this, T cells were cultured and expanded as before for 7 days in control RPMI, and then restimulated for 6 hours in either control RPMI or in TIFM. T cells restimulated in TIFM had small but significant reductions in IFNγ^+^TNF^+^ cells and Granzyme B and IL-2 expression (Figure 1H-J). Collectively, these data indicate that exposure to tumor nutrient stress induces acute effects on T cell function.

### Arginine depletion in TIFM prevents T cell proliferation

To understand how tumor nutrient stress alters both T cell expansion and function, we next sought to determine the critical metabolic constituents of TIFM that drove a dysfunctional phenotype. Mass spectrometry data from tumor interstitial fluid and plasma in PDAC tumor-bearing mice revealed many depleted metabolites in the tumor interstitial fluid when compared to plasma (Figure S2A). To determine whether a reduction in any of these metabolites was driving dysfunction in CD8^+^ T cells in our TIFM system, we performed ‘add-back’ experiments, activating CD8^+^ T cells in RPMI, then transferring the activated CD8^+^ T cells into TIFM, or into TIFM supplemented with individual metabolites at RPMI levels (Figure 2A)^24^. Analysis of T cell expansion revealed that supplementation of arginine, but no other depleted metabolite tested, could promote T cell expansion in TIFM (Figure 2B). Arginine is one of the most limited nutrients in the murine PDAC tumor microenvironment, with a 20-50-fold decrease compared to murine plasma, and a roughly 500-fold decrease compared to RPMI (Figure S2A)^24^. Further, arginine has been shown to be critical for T cell proliferation^27,28^. Arginine supplementation of TIFM also resulted in lower PD-1 expression than TIFM alone, but still, cells cultured in TIFM supplemented with arginine still had significantly higher levels of PD-1 than RPMI-controls (Figure 2C). Additionally, arginine supplementation of TIFM failed to rescue any decreases in cytokine expression in restimulated T cells (Figures 2D-2F). Since arginine supplementation can rescue T cell proliferation but not effector function, these data suggest that TIFM-induced effector dysfunction is uncoupled from the antiproliferative effects of TIFM, and an alternative, TIFM-specific metabolite must be driving these changes to T cell function.

**Figure 2:**
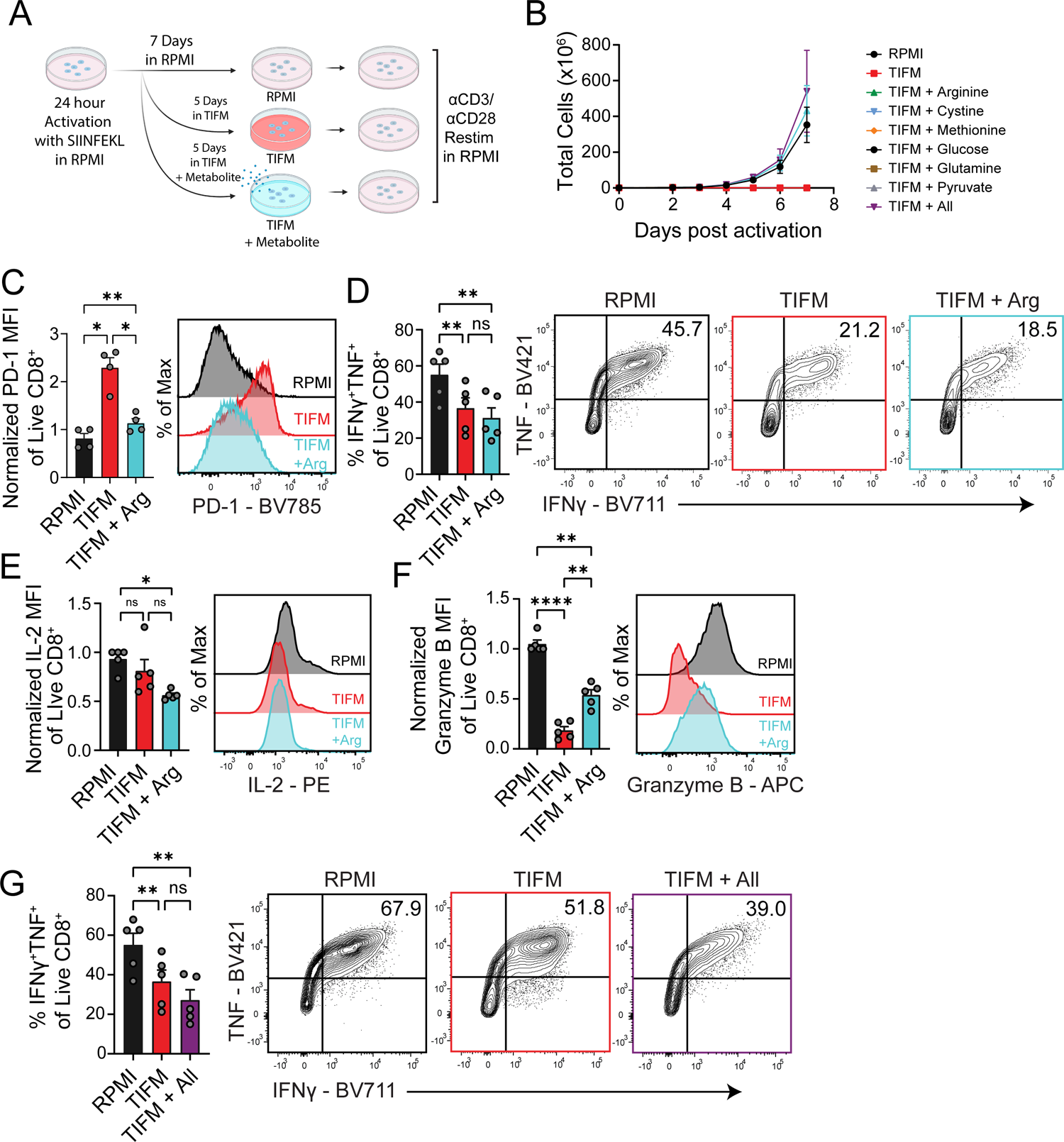
TIFM-derived arginine starvation impairs CD8^+^ T cell proliferation **B)** Splenocytes from OT-I mice were activated with 250ng/mL SIINFEKL peptide for 24 hours in RPMI supplemented with 50U/mL IL-2. Cells were cultured for an additional 24 hours in complete RPMI supplemented with IL-2 and transferred into Tumor Interstitial Fluid Like Medium (TIFM) with IL-2 for 5 days, or TIFM with individual metabolites added back at identical concentrations to complete RPMI. “All” metabolites indicates when each of arginine, cystine, methionine, glucose, glutamine, and pyruvate were all added back to RPMI-levels. Control cells were cultured in RPMI throughout. After 7 days, all cells were restimulated in RPMI with 3µg plate-bound αCD3 and 1µg soluble αCD28 antibody for 6 hours. **B)** Cell expansion from cell culture conditions described in A). Plot shows Total cell number (x10^6^) ± SEM (n=5 from 2 independent experiments). **C)** Expression of PD-1 from Day 7 OT-I CD8^+^ T cells from indicated culture conditions and quantification of normalized PD-1 MFI ± SEM from two independent experiments is shown on the left (n=4/group). **D)** Day 7 OT-I CD8^+^ T cells were analyzed for IFNγ and TNF co-expression after 6 hours αCD3/αCD28 restimulation. Data quantified on the left show % IFNγ^+^TNF^+^ events among Live CD8^+^ T cells ± SEM from three independent experiments (n=5/group). **E-F)** Day 7 OT-I T cells were analyzed for expression of **E)** Granzyme B or **F)** IL-2 after 6 hours of αCD3/αCD28 restimulation. Data quantified on the left show normalized MFI ± SEM from three independent experiments (n=5/group). **G)** Day 7 OT-I CD8^+^ T cells from indicated culture conditions were analyzed for IFNγ and TNF co-expression after 6 hours αCD3/αCD28 restimulation. Data quantified on the left show % IFNγ^+^TNF^+^ events among Live CD8^+^ T cells ± SEM from three independent experiments (n=5/group). Statistical analysis for all indicated comparisons in this figure were calculated using a one-way ANOVA with Tukey’s multiple comparisons test. For indicated comparisons, * = p<0.05, ** = p <0.01, and **** = p<0.0001.

### Phosphoethanolamine, a TIFM-enriched metabolite, induces profound T cell dysfunction

Notably, even restoration of all the tested TIF-depleted metabolites to TIFM to RPMI levels could not restore effector function as measured by the percentage IFNγ^+^TNF^+^ cells (Figure 2G), suggesting that the dysfunctional effects of TIFM culture may not be induced through metabolite depletion, but potentially due to the presence of a deleterious metabolite elevated in TIFM. Thus, we sought to determine the immunologic effects of metabolites enriched in TIFM on effector T cells. To do this, we activated T cells in RPMI as before, but, in addition to culture in TIFM, we instead added TIF-enriched metabolites identified by metabolomics on TIF and plasma (Figure S2A) to RPMI at TIFM-equivalent concentrations (Figures 3A). Our initial screen identified phosphoethanolamine (pEtn), a metabolite which forms the head group of the phospholipid phosphatidylethanolamine (PE), as a potential inhibitory metabolite present in TIFM (Figure 3B-D, blue bars). We found that the effect of pEtn was comparable to the effect of all the tested TIFM-enriched metabolites pooled together, suggesting pEtn as a key regulator of T cell dysfunction in TIFM (Figure 3B-D). In our experiments, pEtn had no negative effect on T cell expansion (Figure 3E). Further, pEtn addition resulted in elevated levels of co-inhibitory molecule (PD-1, Tim-3, Lag3) expression (Figure 3F-H). Additionally, prolonged cell culture in RPMI with pEtn supplementation resulted in a dysfunctional effector phenotype, characterized by decreased IFNγTNF co-expression, as well as IL-2 and Granzyme B production after restimulation with anti-CD3 and anti-CD28 in RPMI (Figures 3I-3K). Next, we tested whether culture in RPMI supplemented with pEtn had a similar time dependent effect on T cell dysfunction to that of TIFM nutrient stress. To test this, OT-I T cells were activated for 24 hours with SIINFEKL peptide, then expanded in RPMI, or transferred into RPMI supplemented with pEtn for 1, 3, or 5, days prior to analysis on day 7 (Figure 3L).

**Figure 3:**
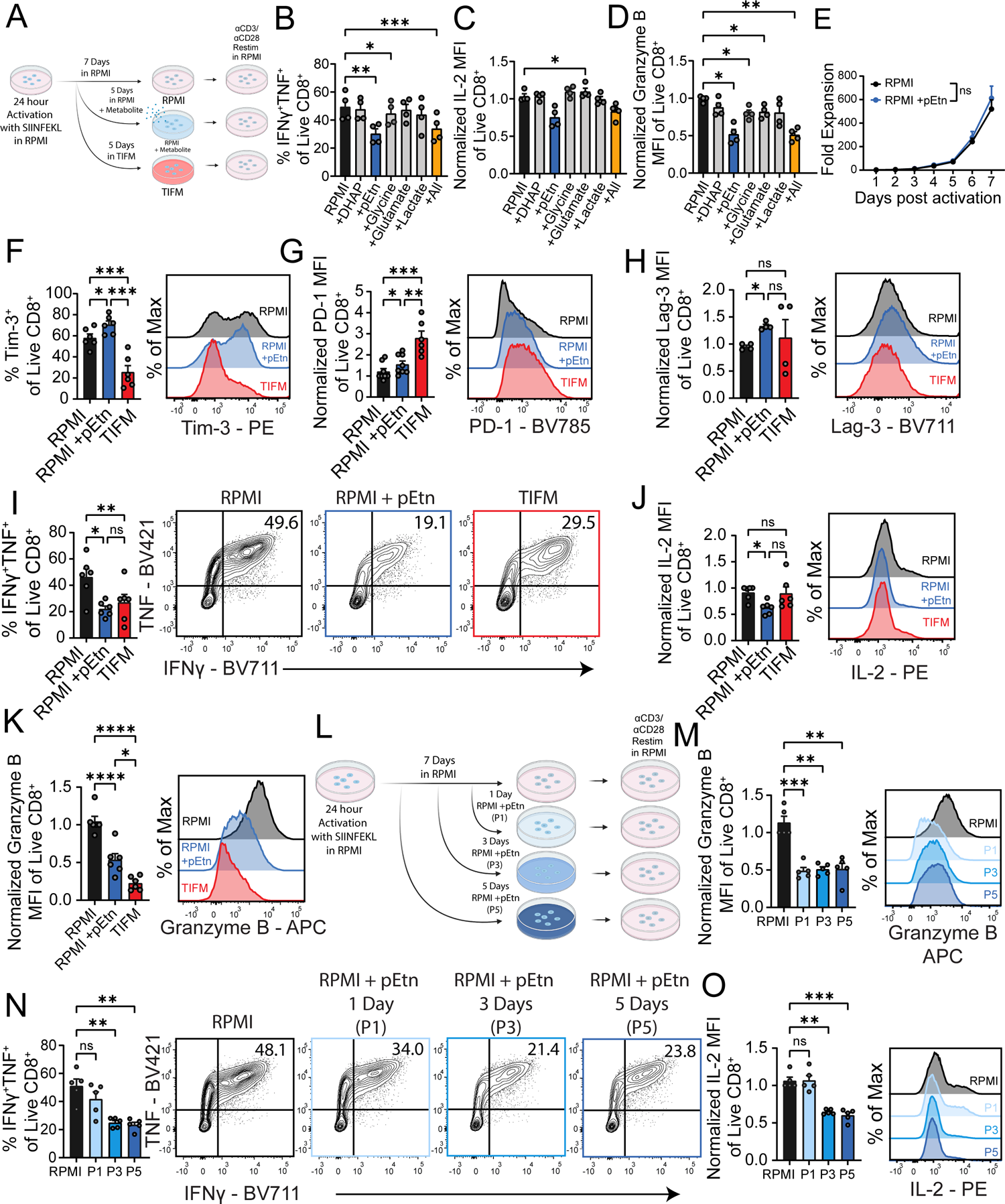
Phosphoethanolamine, a metabolite enriched in tumor interstitial fluid, induces profound T cell dysfunction. **A)** Splenocytes from OT-I mice were activated with 250ng/mL SIINFEKL peptide for 24 hours in RPMI supplemented with 50U/mL IL-2. Cells were cultured for an additional 24 hours in complete RPMI supplemented with IL-2 and transferred into Tumor Interstitial Fluid Like Medium (TIFM) with IL-2 for 5 days, or RPMI with individual metabolites added back at identical concentrations to TIFM. Control cells were cultured in RPMI throughout. After 7 days, all cells were restimulated in RPMI with 3µg plate-bound αCD3 and 1µg soluble αCD28 antibody for 6 hours. **B** Day 7 OT-I CD8^+^ T cells were analyzed for IFNγ and TNF co-expression after 6 hours αCD3/αCD28 restimulation. Data quantified show % IFNγ^+^TNF^+^ events among Live CD8^+^ T cells ± SEM from two independent experiments (n=4/group). **C-D)** Day 7 OT-I T cells were analyzed for expression of **C)** Granzyme B or **D)** IL-2 after 6 hours of αCD3/αCD28 restimulation. Data quantified show normalized MFI ± SEM from two independent experiments (n=4/group). **E)** Fold expansion of cells cultured in A), (n=6 from 3 independent experiments) **F)** Expression of Tim-3 from Day 7 OT-I CD8^+^ T cells from indicated culture conditions and quantification of % Tim-3^+^ ± SEM on the left from three independent experiments is shown on the left (n=6/group). **G)** Expression of PD-1 from Day 7 OT-I CD8^+^ T cells from indicated culture conditions and quantification of normalized PD-1 MFI ± SEM on the left from three independent experiments is shown on the left (n=6/group). **H)** Expression of Lag-3 from Day 7 OT-I CD8^+^ T cells from indicated culture conditions and quantification of normalized Lag-3 MFI ± SEM on left from two independent experiments is shown on the left (n=4/group). **I)** Day 7 OT-I CD8^+^ T cells were analyzed for IFNγ and TNF co-expression after 6 hours αCD3/αCD28 antibody restimulation. Data quantified on the left show % IFNγ^+^TNF^+^ events among Live CD8^+^ T cells ± SEM from three independent experiments (n=6/group). **J-K)** Day 7 OT-I T cells were analyzed for expression of **J)** Granzyme B or **K)** IL-2 after 6 hours of αCD3/αCD28 restimulation. Data quantified show normalized MFI ± SEM from three independent experiments (n=6/group). **L)** Splenocytes from OT-I mice were activated with 250ng/mL SIINFEKL peptide for 24 hours in RPMI supplemented with 50U/mL IL-2. Cells were cultured for an additional 24 hours in complete RPMI supplemented with IL-2 and transferred into RPMI with pEtn added back at identical concentrations to TIFM for 1, 3, or 5 days (P1, P3, P5). Control cells were cultured in RPMI throughout. After 7 days, all cells were restimulated in RPMI with 3µg plate-bound αCD3 and 1µg soluble αCD28 antibody for 6 hours. **M-O)** Day 7 OT-I T cells were analyzed for expression of **M)** Granzyme B **N)** IFN^+^TNF^+^ coexpression or **O)** IL-2 after 6 hours of αCD3/αCD28 restimulation. Data quantified show normalized MFI ± SEM from three independent experiments (n=5/group). Statistical significance for all indicated comparisons in this figure was calculated using a one-way ANOVA with Tukey’s multiple comparisons test. **E)** is the only exception to this, as the indicated comparison is a 2-way ANOVA. For all indicated comparisons, * = p<0.05, ** = p <0.01, *** = p<0.001, and **** = p<0.0001.

Similar to TIFM exposure, cells cultured for 3 or more days in RPMI supplemented with pEtn had the most striking differences in IFNγ, TNF, and IL-2 expression after restimulation when compared to control RPMI-cultured cells, while Granzyme B expression was reduced after just 1 day of pEtn of exposure (Figures 3M-O). These experiments suggested that pEtn, an oncometabolite enriched in the TME, can promote a state of T cell dysfunction on its own.

### The ethanolamine arm of the Kennedy pathway can induce T cell dysfunction

We next wanted to explore how common pEtn exposure may be to tumor-infiltrating T cells in other malignancies. Mass spectrometric quantification confirmed pEtn was enriched in tumor interstitial fluid from other murine tumor models including B16 melanoma, and BRCA^-/-^ p53^+/-^ breast tumors. (Figure 4A). Thus, elevated pEtn is a shared feature of the nutrient environment of multiple tumor types. The finding that elevated pEtn is a common feature of many tumor types motivated us to understand how exposure to extracellular pEtn triggers T cell dysfunction. pEtn is formed from the action of ethanolamine kinase, which phosphorylates ethanolamine (Etn) (Figure 4B). Accumulation of pEtn in TIF is consistent with recent studies that TME conditions repress activity of *Pcyt2* in cancer cells^29^, the enzyme which converts pEtn into CDP-Ethanolamine (CDP-Etn) in the Kennedy pathway of phosphatidylethanolamine (PE) membrane lipid synthesis (Figure 4B). This allows cancer cells to accumulate pEtn enabling survival of TME stress^29^. The Kennedy pathway consists of two arms: 1) an ethanolamine arm to generate PE (Figure 4B, left), and 2) a choline arm to generate phosphatidylcholine (PC) (Figure 4B, right). PE and PC are the two most abundant phospholipids in mammalian cells.

**Figure 4:**
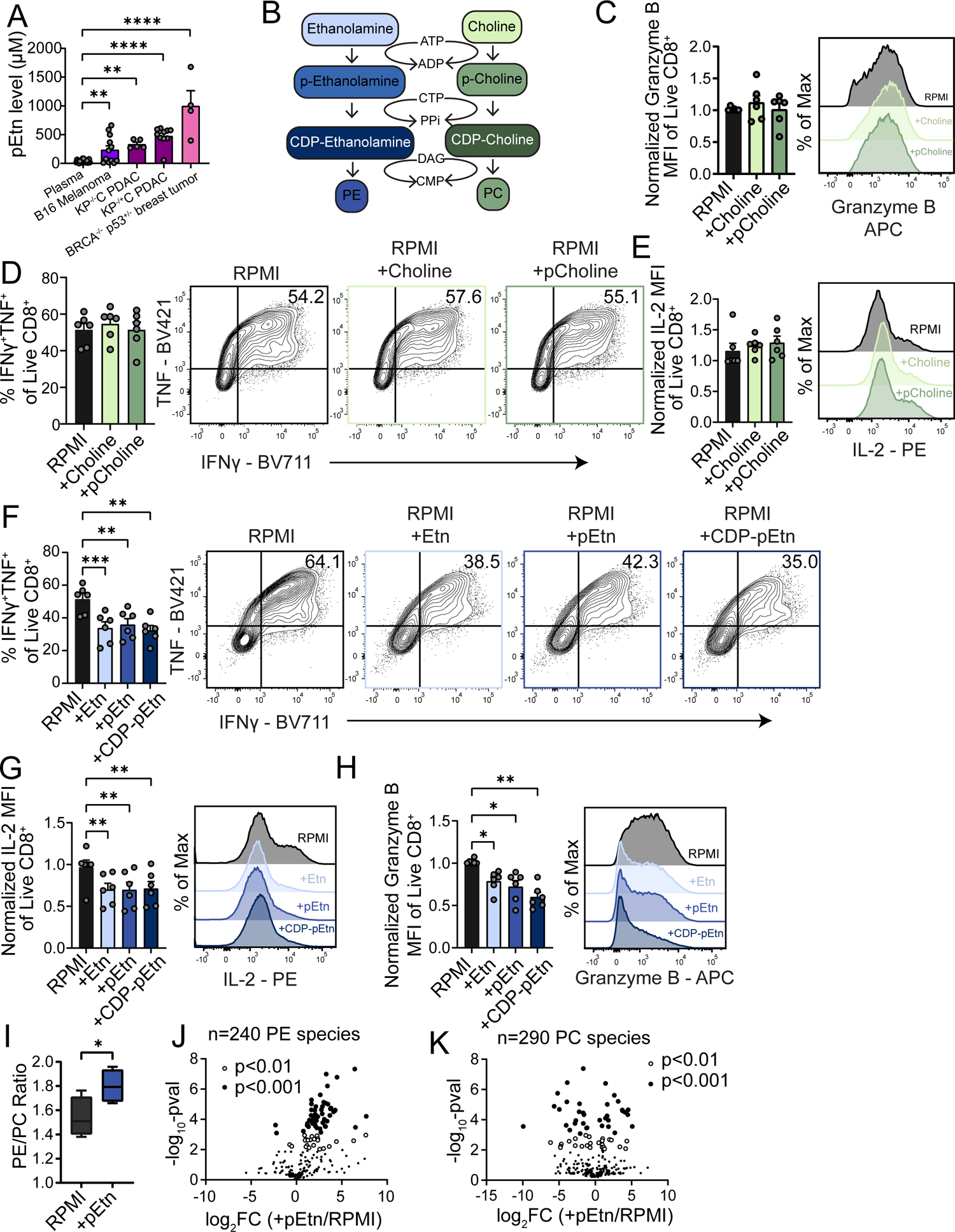
The ethanolamine arm of the Kennedy pathway induces T cell dysfunction **A)** Bar graph showing the pEtn level measured in murine plasma, or in tumor interstitial fluids of listed tumor types: B16 melanoma, KP^-/-^C PDAC, KP^-/+^C PDAC, BRCA ^-/-^ p53^+/-^ breast tumor. Statistical significance for indicated comparisons determined by ordinary one-way ANOVA. **B)** Illustration of the Kennedy pathway, featuring the ethanolamine (blue, left) and choline (green, right) arms of the pathway. **C-E)** OT-1 splenocytes were activated with SIINFEKL for 24 hours, and then cultured in complete RPMI for 24 more hours. Then, cells were transferred into RPMI supplemented with choline or phosphocholine or kept in control RPMI for 5 days. On day 7, cells were restimulated with αCD3/αCD28 antibodies for 6 hours, and **C)** Granzyme B expression, **D)** IFNγ and TNF co-expression, and **E)** IL-2 protein expression was measured via flow cytometry. Data quantified show normalized MFI ± SEM or %IFN^+^TNF^+^ events on the left of each panel from three independent experiments (n=6/group). **F-H)** OT-1 splenocytes were activated with SIINFEKL for 24 hours, and then cultured in complete RPMI for 24 more hours. Then, cells were transferred into RPMI supplemented with ethanolamine (Etn), phosphoethanolamine (pEtn) or CDP-Ethanolamine (CDP-Etn) or kept in control RPMI for 5 days. On day 7, cells were restimulated with αCD3/αCD28 antibodies for 6 hours, and **F)** IFNγ and TNF co-expression **G)** IL-2 expression, and **H)** Granzyme B expression was measured via flow cytometry. Data quantified show normalized MFI ± SEM of indicated target or % IFN^+^TNF^+^ ± SEM on the left of each panel from three independent experiments (n=6/group). Statistical Significance for all indicated comparisons in **C-H)** were calculated using a one-way ANOVA with Tukey’s multiple comparisons test. **I-K)** Lipidomics analysis of activated OT-I T cells cultured in either RPMI or RPMI supplemented with pEtn. **I)** Ratio of phosphatidylethanolamine (PE) to Phosphatidylcholine (PC). Fold change of **J)** PE species and **K)** PC species in pEtn treated OT-I T cells. For all indicated comparisons, * = p<0.05, ** = p <0.01, and *** = p<0.001.

Choline and phosphocholine concentrations were increased in murine PDAC TIF compared to plasma (Figure S3A), so we first asked whether metabolites in the choline arm Kennedy pathway metabolites also drove T cell dysfunction. To test this, we supplemented RPMI with choline and phosphocholine (pCholine) at TIFM-equivalent levels and cultured activated CD8^+^ T cells in this media as with other metabolite additions before (Figure 3A).

Neither choline nor phosphocholine supplementation induced any changes in Granzyme B, IFNγ, TNF or IL-2 cytokine expression after αCD3/αCD28 restimulation when compared to control RPMI cultured cells (Figures 4C-4E). We next tested whether supplementation with Kennedy pathway metabolites in the ethanolamine arm could drive dysfunction. Both the precursor of pEtn, Etn, and its enzymatic product, CDP-Etn, reduced IFNγ and TNF co-expression as well as IL-2 and Granzyme B production after restimulation with αCD3/αCD28 (Figures 4F-4H).

Given the overabundance of pEtn in TIFM relative to plasma or RPMI, we hypothesized that there may be an altered balance toward PE lipid species in CD8^+^ T cells cultured in high pEtn conditions. To test this, we cultured activated OT-I CD8^+^ T cells in RPMI or RPMI supplemented with pEtn at TIFM-equivalent concentrations for 5 days and conducted lipidomics on each group of cells. As expected, pEtn-treated CD8^+^ T cells had a heightened PE/PC ratio (Figure 4I). This effect was largely driven by an overwhelming increase in PE species abundance, in which 95% (79 of the 83) of the differentially abundant PE Species (as defined by Log2FC >2 and a p-value <0.01) were increased in pEtn treated cells compared to control T cells (Figure 4J). PC species were more evenly affected, with only 47% (29 of the 61) differentially abundant PC species being increased in pEtn treated cells compared to controls, while the other 53% were decreased (Figure 4K). These data confirm that pEtn availability drives anabolic activity in the PE arm of the Kennedy pathway of CD8^+^ T cells. Further, increased availability of these ethanolamine species drives a dysfunctional phenotype in CD8^+^ T cells.

### Phosphoethanolamine-induced T cell dysfunction is not canonical exhaustion but alters TCR signal transduction

We next wanted to understand the underlying cellular consequences of pEtn exposure in T cells. Global metabolomics profiling showed pEtn supplementation resulted in significant reprogramming of carbohydrate metabolism and confirmed accumulation of pEtn and its downstream substrates (Figures S4A and S4B). Most TCA cycle metabolites, intermediates of glycolysis and glutaminolysis, and lipid metabolites were suppressed (Figure S4A). Moreover, sphingomyelin was elevated, suggesting pEtn-derived PE was being further converted into downstream metabolites, depleting ceramides as a consequence (Figure S4B). We also performed RNAseq on pEtn expanded T cells, which identified several key immunologic genes altered by pEtn treatment (Figure S4C). However, there was no notable differentiation pattern associated with pEtn treatment as determined through enrichment analysis of T cell differentiation gene sets (Figure S4D), suggesting pEtn alone was not driving differentiation into canonical CD8^+^ T cell exhaustion. Indeed, consistent with the notion that pEtn exposure does not trigger canonical exhaustion, when pEtn-expanded T cells were restimulated chemically with 13-phorbol-12-myriostol-acetate (PMA) and ionomycin, circumventing the need for proximal TCR signaling, there was no observable functional defect in effector cytokine synthesis (Figures S4E-G). Thus, exposure to pEtn did not change the intrinsic capacity of the CD8^+^ T cells to make TNF, IFN, and IL-2, or Granzyme B, but rather dampened their ability to respond to T Cell Receptor (TCR) stimulation.

### Phosphoethanolamine inhibits TCR signal propagation through depletion of DAG

We next wanted to understand the molecular mechanisms through which pEtn exposure impaired the response of the T cells to TCR stimulation. Therefore, we examined the TCR signaling cascade in T cells exposed to TIF-equivalent levels of pEtn. Interestingly, restimulation with anti-CD3 and anti-CD28 showed proximal TCR signaling was retained in pEtn-exposed T cells, including phosphorylation of Lck, ZAP70, and PLCγ (Figures 5A-5D). However, downstream signaling through the MAPK/ERK and Akt signaling pathways was significantly repressed in pEtn-expanded T cells (Figures 5A, 5E, and 5F). Stimulation of T cells on stimulatory lipid bilayers showed that TCR microclustering could still be induced in pEtn treated cells, although TCRs did not cluster tightly resulting in a larger, more diffuse supramolecular activation cluster (SMAC) (Figure 5G), consistent with inhibited downstream signaling. The signaling pathways downstream of TCR engagement that are repressed in pEtn-exposed T cells require Phospholiopase C, gamma 1 (PLCγ) production of the second messenger metabolites: diacylglycerol (DAG) and IP3^30^. The Kennedy pathway reaction that generates PE from CDP-Etn utilizes DAG as a substrate (Figure 4B). Thus, we hypothesized that heightened flux through the Kennedy pathway through pEtn supplementation may deplete this critical second messenger, inhibiting the amplification of the TCR signal through DAG depletion. Indeed, flow cytometry measurement of DAG on cells cultured in RPMI supplemented with pEtn confirmed that pEtn supplementation depleted the steady state levels of DAG in T cells (Figure 5H). This effect was specific to DAG, as calcium flux induced by TCR stimulation (IP3) was unaltered in pEtn-exposed T cells (Figure 5I). PMA, commonly used to chemically activate T cells, functions as an analog of DAG. Given this, we hypothesized that PMA supplementation could restore cytokine production in the DAG-depleted pEtn-treated CD8^+^ T cells upon αCD3 restimulation.

**Figure 5:**
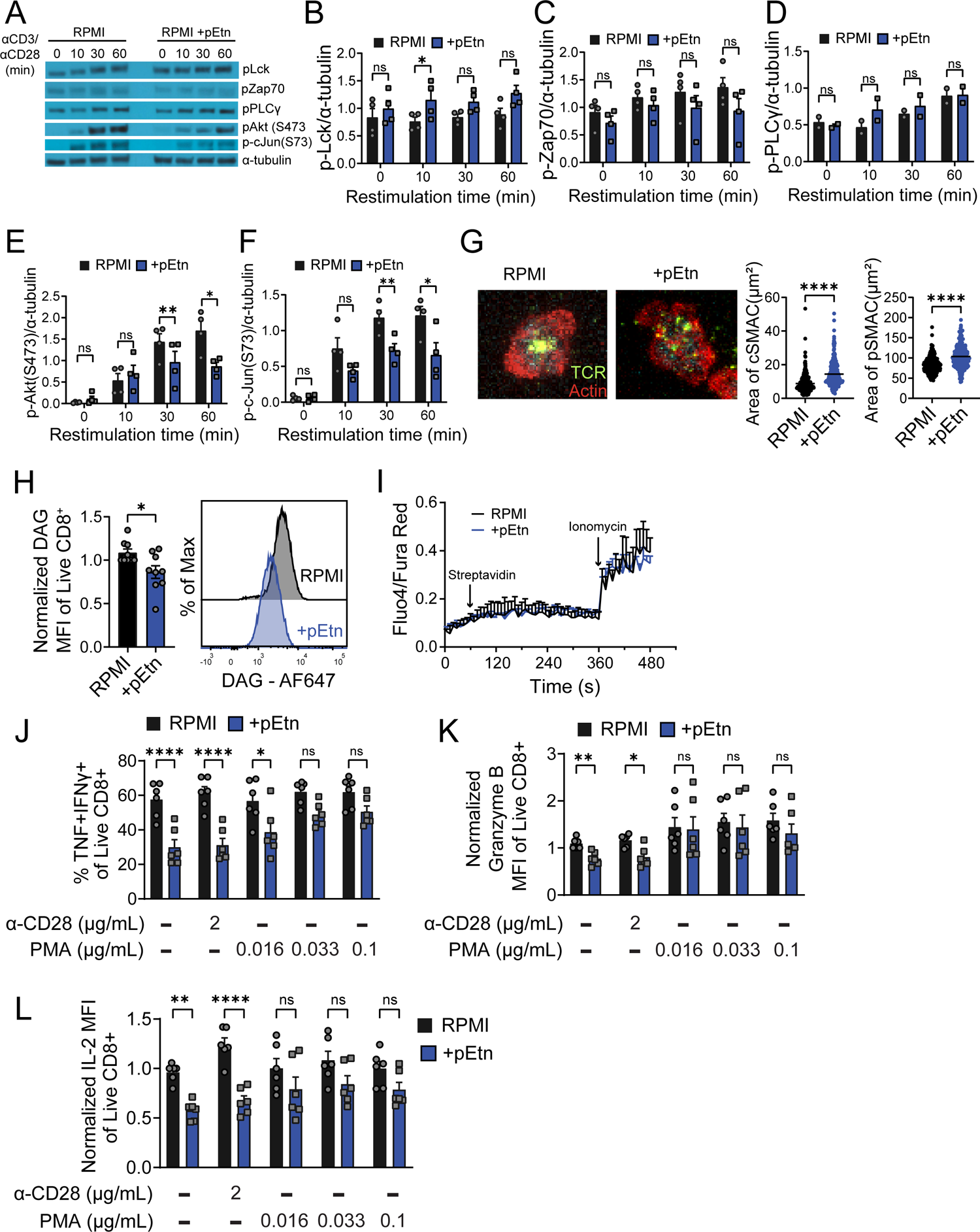
Phosphoethanolamine depletes diacylglycerol (DAG) from CD8^+^ T cells and impairs their function. **(A-F)** OT-I T cells activated for 24 hours were cultured in RPMI for 2 days and then in either RPMI or RPMI+pEtn for 5 days as described in Figure 3A. On day 7, cells were restimulated with αCD3/αCD28 for 10, 30, or 60 minutes, or left unstimulated. Western blotting was then conducted on lysates from stimulated cells. Representative blots for all proteins are shown in **A)** with quantitation from four independent experiments for **B)** p-Lck **C)** p-Zap70 **D)** p-Plcγ **E)** p-Akt (Ser473) and **F)** p-c-Jun (n=4/group). Statistical significance for all indicated comparisons in **A-F)** was calculated using a paired t-test. **G)** OT-I T-cells were cultured as described in **A-F**), on Day 7, synapse formation was monitored on stimulatory lipid bilayers cSMAC is indicated by TCR (Green) clustering and pSMAC area is indicated by actin (red). Tubulated data of cSMAC and pSMAC area of the cells are shown on the right. Statistical significance for indicated comparisons was calculated using an unpaired t-test. **H)** DAG levels in OT-I T cells that were cultured as described **in A-F).** Normalized DAG MFI is quantified on the left from five independent experiments and statistical significance was calculated using a paired t-test (n=9/group). **(I)** Intracellular calcium flux analysis with Fluo4/Fura Red on OT-I T cells cultured as in **A-F)** in response to α-CD3 signal (streptavidin) and ionomycin (n=2 from two independent experiments). **J-L)** OT-I T cells cultured as in **A-F)** were restimulated with αCD3 and either αCD28 or PMA at the indicated concentration for 6 hours in RPMI and analyzed for cytokine production. **J)** IFNγ and TNF coexpression, **K)** Granzyme B expression, and **L)** IL-2 expression were measured via flow cytometry from three independent experiments (n=6/group). Statistical significance of the indicated comparisons was calculated using a two-way ANOVA with Sidak’s multiple comparisons test. * = p<0.05, ** = p <0.01, and **** = p<0.0001.

Indeed, PMA supplementation restored cytokine production of pEtn-treated T cells in a dose-dependent manner, with no effects observed in control T cells (Figure 5J-L). Thus, pEtn supplementation drives heightened flux through the ethanolamine arm of the Kennedy pathway, depleting DAG, and blunting TCR signal transduction during reactivation.

### Prevention of phosphoethanolamine accumulation in tumors promotes tumor-infiltrating CD8^+^ T cell function

Finally, we wanted to confirm the importance of these pathways *in vivo* in tumor-infiltrating T cells. In order to study antigen specificity in an established tumor model that induces strong T cell dysfunction, we elected to use B16 melanoma, which, importantly, accumulates pEtn (Figure 4A). Mice were inoculated with B16 melanoma and received an adoptive transfer of Pmel-1 (gp100-specific) congenically mismatched T cells (Figure 6A). The transferred T cells exhibited DAG depletion when isolated from tumors compared to lymph nodes (Figure 6B). These data support the idea that the tumor microenvironment drives depletion of DAG in CD8^+^ T cells. Notably, consistent with our previous data suggesting pEtn did not induce canonical exhaustion, DAG depletion did not correlate with terminal differentiation as marked by PD1 and Tim3 co-expression in either the endogenous (Figure 6C) or transferred tumor infiltrating T cells (Figure S5A). Finally, we asked whether we could generate a tumor microenvironment with decreased pEtn accumulation and measure the effect pEtn depletion would have on the intratumoral T cell phenotype. Overexpression of Pcyt2 in HeLa cells was previously reported to reduce intratumoral pEtn concentrations^29^. We generated a B16 melanoma line that stably overexpresses *Pcyt2* (Figure S5B) to see if reducing TIF pEtn levels could improve tumor infiltrating lymphocyte function. As expected, Pcyt2-overexpressing cells had reduced levels of pEtn abundance in the cells (Figure 6D) and the cell culture supernatant (Figure 6E) compared to B16-empty vector (B16-EV) controls. Analysis of the interstitial fluid of implanted tumors showed pEtn was significantly decreased in TIF from B16-Pcyt2 compared to TIF from B16-EV, although a sizeable proportion of pEtn remained (Figure 6F). While B16-*Pcyt2* tumors grow at comparable rates when implanted into in Rag2^-/-^ mice (Figure 6G), they grow significantly slower when implanted into wild type C57/BL6 mice (Figure 6H). These data indicate that the decreased tumor growth in wild-type C57/BL6 mice is due to antitumor adaptive immunity. To better characterize the T cell infiltrate, WT B6 mice implanted with B16-Wt or B16-Pcyt2 tumors and given pmel T cell transfer were analyzed 10 days after T cell transfer (Figure 6I). Tumor area of B16-Pcyt2 tumors was two-fold smaller than B16-EV control tumors at the day 10 harvest (Figure 6J). Functionally, Tumor-infiltrating Pmel T cells from B16-*Pcyt2* were more capable of producing TNF and IL-2 after gp100 restimulation (Figure 6K-L). Additionally, tumor-infiltrating Pmel T cells from B16-Pcyt2 tumors had a 2.5-fold increase in DAG levels when compared to B16-EV controls (Figure 6M). Further, the expression of PD-1 inhibitory receptor on the surface of transferred Pmel T cells was lower in cells harvested from B16-Pcyt2 tumors than B16-EV controls (Figure 6N). Collectively, these data show that a reduction in pEtn accumulation in tumor microenvironments improves tumor-specific CD8^+^ T cell functions antitumor adaptive immunity. Thus, pEtn represents a novel oncometabolite produced and secreted at high levels into the TME by cancer cells that induces a distinct form of T cell dysfunction induced through pathologic flux of the Kennedy pathway, which depletes the essential TCR signaling second messenger DAG.

**Figure 6:**
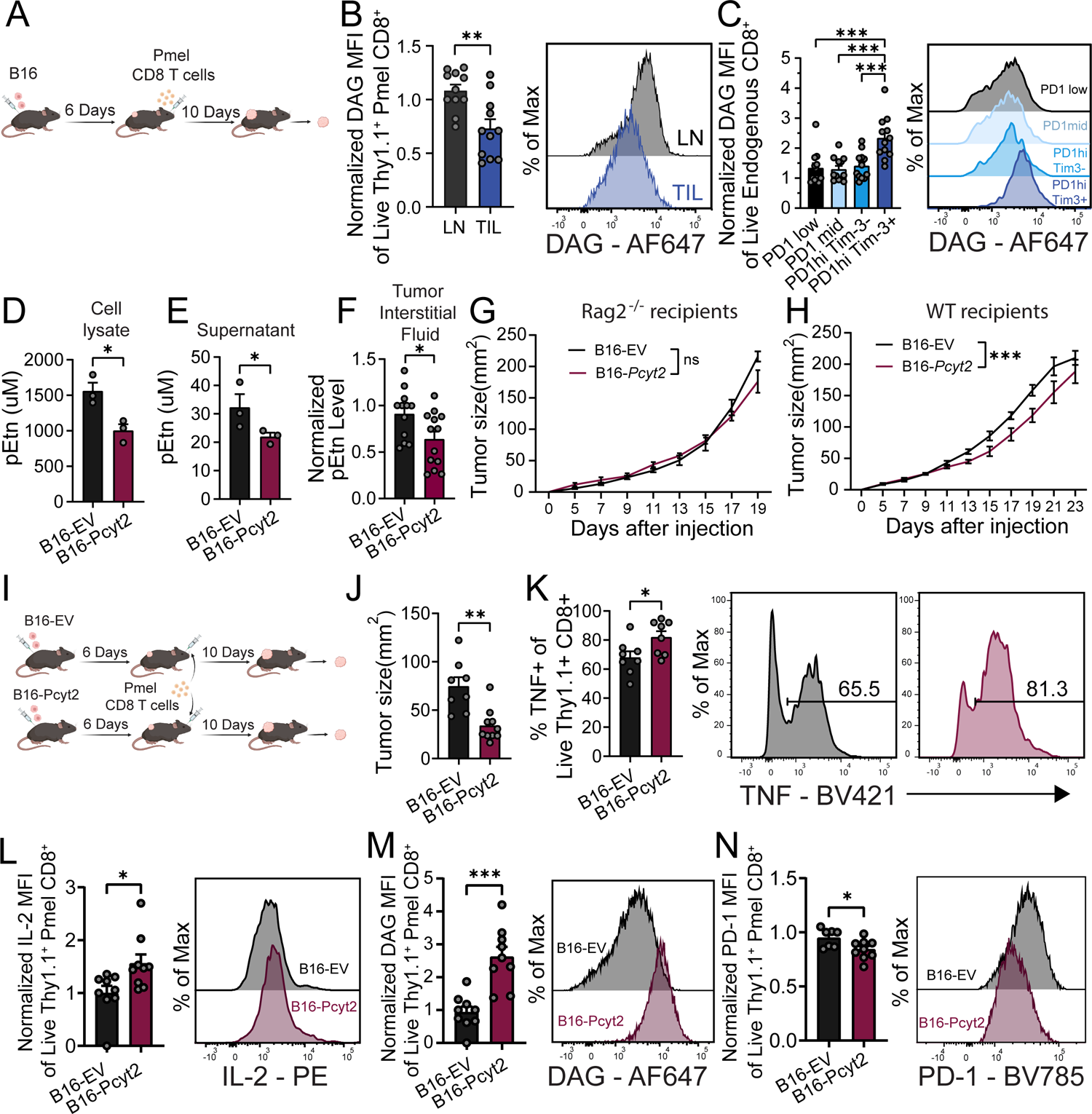
Depletion of phosphoethanolamine in the tumor microenvironment promotes increased antitumor immunity. **A)** C57BL/6J mice were injected subdermally with B16 melanoma cells. 6 days after tumor cell injection, Pmel T cells were transferred retro-orbitally into tumor-bearing mice. 10 days post T cell transfer, tumors and draining lymph nodes were harvested and analyzed. **B)** DAG levels from transferred T cells within lymph node (LN) and Tumor (TIL). Normalized DAG MFI quantified on the left is from 2 independent experiments (n=11/group). Statistical significance for indicated comparison calculated using an unpaired t-test. **C)** DAG levels of endogenous CD8^+^ TILs with gated on PD1low, PD1mid, PD1hi Tim3-, and PD1hiTim3^+^. Normalized DAG MFI quantified on the left is from 2 independent experiments (n=12/group). Statistical significance of indicated comparisons was calculated using a one-way ANOVA with Tukey’s multiple comparisons test. **D-E)** B16 melanoma cells were transduced with empty vector (B16-EV) or Pcyt2 overexpression constructs (B16-Pcyt2). Cells were cultured in RPMI, and levels of pEtn were measured in **D)** cell lysates and **E)** Supernatant. Data are from three independent experiments (n=3/group) **F)** B16-EV or B16-Pcyt2 cells were injected subdermally into C57BL/6J mice, and pEtn levels were measured from the tumor interstitial fluid at endpoint. Data quantified in bar plot are from two independent experiments (n=12-13/group) Statistical significance for indicated comparisons in **D-F)** were calculated using an unpaired t-test. **G-H)** B16-EV or B16-Pcyt2 melanoma cells were injected subdermally into **G)** Rag2^-/-^ or **H)** WT C57BL/6J recipients. Tumor growth curve is shown ± SEM. Statistical significance for indicated comparisons was calculated using repeated measures ANOVA. **I)** C57BL/6J mice were injected subdermally with B16-EV or B16-Pcyt2 melanoma cells. 6 days after tumor cell injection, Pmel T cells were transferred retro-orbitally into tumor-bearing mice. 10 days post T cell transfer, tumors were harvested and analyzed. **J)** Tumor size from B16-EV and B16-Pcyt2 tumors on day 16 in experiment described in **I)**. **K-L)** tumor infiltrating pmel T cells were restimulated with gp100 peptide for 6 hours and analyzed for **K)** TNF expression and **L)** IL-2 expression TNF^+^ and normalized IL-2 MFI quantified on the right from 2 independent experiments (n=8-9 group). **M)** DAG levels tumor infiltrating transferred pmel T cell by flow cytometry. Data quantified on left are from one independent experiment (n=8-9/group) **N)** PD-1 expression of tumor infiltrating transferred pmel T cells by flow cytometry. Data quantified on the left are from one independent experiment (n=7-9/group). Statistical significance from all indicated comparisons in J-N were calculated using an unpaired t-test. * = p<0.05, ** = p <0.01, *** = p<0.001, and **** = p<0.0001.

## Discussion

Tumors use multiple mechanisms to induce immune dysfunction. Better understanding of these mechanisms led to therapies designed to reinvigorate adaptive immunity, and therapies targeting co-inhibitory signals have achieved success. However, the relatively low response rates of immunotherapies like checkpoint blockade highlight the multifactorial nature of tumor-induced T cell dysfunction. We now appreciate that tumor nutrient stress also has significant impacts on many anti-tumor immune responses. In this study, we employed a media formulation with nutrient levels equivalent to PDAC tumor interstitial fluid to isolate the specific effects of tumor nutrient stress on T cell function. In doing so, we found that tumor nutrient stress is sufficient to trigger CD8^+^ T cell dysfunction and confirmed the critical importance of arginine availability for T cell expansion. We also identified phosphoethanolamine as a novel oncometabolite that induces profound CD8^+^ T cell dysfunction.

Activated T cells are exquisitely sensitive to their surrounding nutrient conditions, differentiating and performing differently depending on the nutrients readily available to them^13,31,32^. Here, we build upon previous work examining the metabolism and phenotype of immune cells when grown in media designed to recapitulate physiological conditions. Previous work found T cells proliferate and function in a distinct manner when expanded in media designed to mimic human plasma nutrient levels, due to changes in ionic balances between physiological and standard media^31^. In our work, we have sought to use model systems that recapitulate the pathophysiology of tumors to understand how such conditions impact immune cell biology. We believe our findings highlight the importance of tractable model systems to study immune cell biology under the diverse physiological and pathological environments these cells encounter. Such systems will help propel the study of immunology by providing not just a discovery tool for novel metabolite:cell interactions critical for immunology, but also experimental platforms for the identification of cancer therapeutics and immunologic modulators that target or overcome metabolite:cell interactions that contribute to disease.

Our data identified a unique functional phenotype when CD8^+^ T cells are exposed to tumor nutrient stress as modeled by TIFM. Activated CD8^+^ T cells cultured in TIFM had elevated levels of PD-1, and an impaired expression of cytokines such as Granzyme B, IFNγ, and TNF upon restimulation. Functionally, CD8^+^ T cells cultured in TIFM had impaired tumor cell killing *in vitro* compared to control CD8^+^ T cells. Specifically, this work identified phosphoethanolamine, a Kennedy pathway intermediate, as a key metabolite that accumulates in the tumor microenvironment and plays a potent role in suppressing CD8^+^ T cell function. CD8^+^ T cells cultured in pEtn-containing media or harvested from pEtn-producing tumors had depleted levels of DAG, a critical secondary messenger of the TCR signaling pathway. Importantly, the functional impairment of pEtn-cultured CD8^+^ T cells could be rescued with the addition of PMA, a functional analog of DAG. These findings dovetail with several studies highlighting pEtn production in transformed cells, which point to a shared immunosuppressive mechanism across tumor types. Recent work showed that pEtn production is a hallmark of senescent cells^33^, which are growth arrested cells with dysregulated metabolism commonly found in tumors and premalignant lesions^34^. The secretory phenotype of senescent cells is often considered pro-tumorigenic^35^. Given our data that environmental pEtn induces a hyporesponsive state in CD8^+^ T cells, it is plausible that pEtn production could be a critical step towards tumorigenesis.

Accumulation of pEtn is also common in tumor cells through Pcyt2 silencing, due in part to alterations in the p53 pathway^29^. Other studies show pEtn accumulation in tumors is a result of reduced expression of AGXT2L1, a pEtn phosphorylase and can irreversibly degrade pEtn to acetaldehyde, phosphate and ammonia^36,37^. In T cells, ionic imbalances in the tumor microenvironment trigger metabolic pathways that drive enrichment of Kennedy pathway metabolites, including pEtn, within tumor infiltrating T cells^38^. Our findings suggest that increased abundance of Kennedy pathway metabolites in cancer plays an important role by rendering T cells less capable of tumor control. Thus, we propose pEtn is a novel oncometabolite that tumors and premalignant lesions produce that suppresses CD8^+^ T cell function. This novel mechanism also opens the possibility that therapeutics that target accumulation of pEtn in tumors improve tumor-infiltrating T cell function.

Altogether, our data utilized a novel model system incorporating metabolic parameters of the tumor microenvironment to study the impact of tumor physiology on T cell biology. We both confirmed the importance of known metabolic imbalances but also identified a novel oncometabolite, phosphoethanolamine, that is enriched in many tumor microenvironments, and drives T cell dysfunction in tumors by a novel metabolic mechanism. Thus, our findings support a model in which the nutrient content of the tumor microenvironment, independent from non-metabolic signals, is sufficient to bias T cell differentiation into a dysfunctional fate, highlighting the critical importance of understanding metabolite:immune cell interactions in tumors. Looking forward, we envision TIFM and similar *ex vivo* model systems interrogating tumor physiology will serve as useful platforms to identify how the metabolic microenvironment interfaces with the multiple cell types present in tumors to affect disease progression and therapy response. TIFM may also be used as a critical screening tool to identify targets to improve immune cell function given the metabolic constraints of the tumor microenvironment.

## METHOD DETAILS

### Mice

C57BL/6J mice, OT-I mice, and Pmel-1 mice were obtained from Jackson Laboratories and bred in house. The RAG2 KO mice were obtained from The Jackson Laboratory. These mice when used in experimental procedure were males and females between of 6-8 weeks old. All animal work and protocols in the current study were approved by the University of Pittsburgh Institutional Animal Care and Use Committee, accredited by the AAALAC.

### T cell isolation, activation and culture

Spleen and lymph nodes were isolated from mice, mechanically disrupted, RBC-lysed and then incubated with a biotinylated antibody cocktail consisting of antibodies (BioLegend) to B220, CD11b, CD11c, CD16/32, CD19, CD25, CD105, NK1.1, TCRgd, and CD4. After a wash step, cells were incubated with streptavidin-coated magnetic nanoparticles (BioLegend). After washing, CD8^+^ cells were isolated by applying a magnetic field and removing untouched cells. For peptide activation, the single cell suspension of splenocytes was cultured with peptide (SIINFEKL for OT-I and gp100 for Pmel-1) and IL-2 overnight. For anti-CD3 and anti-CD28 activation, CD8^+^ T cells isolated as above were activated with 3 μg/ml plate-bound anti-CD3, 2 μg/ml soluble anti-CD28 and IL-2 overnight. Activated T cells were cultured in 50U/ml IL2.

### T cell restimulation and staining for flow cytometry

T cells were restimulated with 3 μg/ml plate-bound anti-CD3 and 2 μg/ml soluble anti-CD28 or PMA/Ionomycin in the presence of GolgiPlug protein transport inhibitor (BD Biosciences) for 6 hours. The cells were then stained for cell surface markers for 15min on ice, followed by fixation/permeabilization with fixation/permeabilization solution (BD Biosciences) for 25min in room temperature. The cells were stained for intracellular cytokines at 4 °C overnight and then run on BD LSRFortessa or a Beckman Coulter Life Sciences CytoFLEX.

### xCELLigence real-time killing assay

15,000 B16-OVA cells were seeded onto the E-Plate VIEW 96 (Agilent 300601020) and incubated for a minimum of 5 hours. Subsequently, primary CD8 T cells, isolated from C57BL/6-Tg(TcraTcrb)1100Mjb/J (Strain #:003831, common name: OT1), were introduced to the E-Plate 96 at 1:1 Effector:Target Ratio. Following this, the system was left undisturbed for a minimum of 18 hours to allow the Agilent xCELLigence RTCA DP system to record impedance changes. The RTCA Software Pro was utilized for cytolysis calculations.

### Tumor models, adoptive cell transfer and TIL analysis

B16-F10/B16-veh/B16-Pcyt2 cell lines (100,000 cells intradermally) were injected into the mice on day 0 and followed until the tumors reach 15 mm in any direction. Tumors were measured every other day with digital calipers and tumor size was calculated by LxW for tumor growth curve.

For adoptive transfer, T cells were generated by activating Pmel-1 x Thy1.1 mice with gp100 and expanding the cells for 7 days with IL-2. The cells were then delivered (10^7^) retro-orbitally to mice bearing tumors.

To obtain single-cell suspensions of TILs, tumor bearing mice were sacrificed and tumors were harvested. Excised, the whole tumors were injected using 20G needles with 2mg/mL collagenase type VI, 2U/mL hyluronidase (Dispase), and 10U/mL DNase I (Sigma) in serum-free RPMI and incubated for 30 min at 37 °C. Tumors were then mechanically disrupted between frosted glass slides and filtered to remove particulates, and then vortexed for 2 min.

For restimulation, single-cell suspensions of TILs obtained as above were cultured in culture media with gp100 and GolgiPlug protein transport inhibitor for 6 hours before being stained and run as described above.

### Immunoblotting

Immunoblotting was performed as previously described. Briefly cells were lysed in 1% NP-40 lysis buffer. Cell lysates were then separated by SDS-PAGE using a 4%–12% Bio-Rad gels. Gels were then transferred to a polyvinylidene difluoride membrane and blocked in 5% BSA in Tris-buffered saline 0.1%Tween-20 (TBST). Membrane was incubated overnight at 4°C with primary antibodies diluted in blocking buffer. Membrane was then incubated with secondary antibody (anti-mouse horseradish peroxidase, Jackson ImmunoResearch) in blocking buffer for 1 hour at room temperature and subsequently washed 3 times for 10 minutes with TBST. Protein was visualized by chemiluminescence by using Western Lightning (PerkinElmer).

### Metabolomics analysis

CD8^+^ T cells were activated with peptide and rested in culture media with IL-2, with or without pEtn for 7 days. We flash-froze the cells (10^7^) into pellets. The cells were extracted and analyzed on the Ultrahigh Performance Liquid Chromatography-Tandem Mass Spectrometry (UPLC-MS/MS) platform by Metabolon. The analyzed metabolite dataset comprises a total of 410 biochemicals, 392 compounds of known identity (named biochemicals) and 18 compounds of unknown structural identity (unnamed biochemicals). Following normalization to Bradford protein concentration, log transformation and imputation of missing values, if any, with the minimum observed value for each compound, Welch’s Two Sample t-Test and Matched Pairs t-test were used to identify biochemicals that differed significantly between experimental groups. The data analysis was conducted by Metabolon.

### Calcium flux analysis

Cells were coated with anti-CD3-biotin, stained by fluo-4 and Fura Red and then analyzed on BD LSRFortessa as follows: baseline 1min+ streptavidin 5min+ Ionomycin 2min. Fluo-4/ Fura Red was calculated and plotted against time to show the calcium flux of the cells upon restimulation.

### Imaging of T cell immune synapses on lipid bilayers

Activated CD8^+^ T cells were rested in culture media with IL-2, with or without pEtn for 7 days. Restimulation of the cells on stimulatory lipid bilayer and imaging of the T cell synapse formation were performed as described before^39^. Briefly, Alexa 488 streptavidin and biotinylated TCRβ mAb were sequentially loaded onto the bilayer. Unlabeled poly–His-tagged ICAM-1 produced in a baculovirus system was loaded to facilitate T cell adhesion. The T cells were washed in RPMI without serum or phenyl red for 15 min at 37°C, resuspended in RPMI without serum or phenyl red, and stimulated on a planar lipid bilayer for 10min. The cells were then fixed with 5% paraformaldehyde for 20min, followed by permeabilization with 0.5% Triton X-100 for 5min and blocking with blocking buffer (10% goat serum, 1% BSA and 0.1% Tween 20) for 1h in room temperature. The cells were stained by phalloidin and then imaged with Nikon Spectral A1 Confocal.

### Generation of Pcyt2-overexpressing B16 (B16-Pcyt2) cell line

All cloning was performed using the NEBuilder® HiFi DNA Assembly system. DNA fragments were synthesized using the gBlocks™ platform. The Lentivirus vector pHIV-iRFP720-E2A-Luc was obtained from Addgene. Lentiviruses carrying pcyt2-RFP720 or RFP720 were packaged using the Lenti-X™ 293T Cell Line. Following transduction into B16-F10 melanoma cells, the cells were sorted based on RFP720 fluorescence.

### Untargeted high-resolution LC-HRMS lipidomic analysis

Frozen cell samples were homogenized and then subjected to Folch extraction with 5µL LipidSPLASH deuterated lipid internal standard, and 5µL each deuterated PE/PC UltimateSPLASH internal standards (Avanti Polar Lipids. Alabaster, AL). Analyses were performed by untargeted LC-HRMS. Briefly, Samples were injected via a Thermo Vanquish UHPLC and separated over a reversed phase Thermo Accucore C-18 column (2.1×100mm, 5μm particle size) maintained at 55°C. For the 30 minute LC gradient, the mobile phase consisted of the following: solvent A (50:50 H2O:ACN 10mM ammonium acetate / 0.1% acetic acid) and solvent B (90:10 IPA:ACN 10mM ammonium acetate / 0.1% acetic acid). Initial loading condition is 30% B. The gradient was the following: Over 2 minutes, increase to 43%B, continue increasing to 55%B over 0.1 minutes, to 65%B over 10 minutes, to 85%B over 6 minutes, and finally to 100% over 2 minutes. Hold at 100% for 5 minutes, followed by equilibration at 30%B for 5 minutes. The Thermo IDX tribrid mass spectrometer was operated in both positive and negative ESI mode. A data-dependent MS2 method scanning in Full MS mode from 200 to 1500 m/z at 120,000 resolution with an AGC target of 5e4 for triggering ms2 fragmentation using stepped HCD collision energies at 20, 40, and 60% in the orbitrap at 15,000 resolution. Source ionization settings were 3.5 kV and 2.4kV spray voltage respectively for positive and negative mode. Source gas parameters were 35 sheath gas, 5 auxiliary gas at 300°C, and 1 sweep gas. Calibration was performed prior to analysis using the PierceTM FlexMix Ion Calibration Solutions (Thermo Fisher Scientific). Internal standard peak areas were then extracted manually using Quan Browser (Thermo Fisher Xcalibur ver. 2.7), normalized to weight and internal standard peak area at both the individual lipid and class level, then graphed using GraphPad PRISM (ver 9.0). Untargeted differential comparisons were performed using LipidSearch 4.2 (Thermo Fisher) to generate a ranked list of significant lipid compounds at the class and species-specific levels.

### Phosphoethanolamine extraction and quantification

5 µL of tumor interstitial fluid (TIF), cell culture medium or frozen B16F10 cell pellets (∼2X10^6^ of cells) were extracted with 45 µL of extraction solvent (75:25:0.1 acetonitrile/methanol/formic acid spiked with 13C2 phosphoethanolamine at 20 µM concentration). The calibration curve was prepared by serial dilution of phosphoethanolamine standard to achieve solutions ranging from 500 to 0.01 µM in water. Similar to TIF samples, 5 µL of each calibration curve solution was extracted with 45 µL of extraction solvent. Samples were sonicated for 1 minute, incubated on ice for 20 minutes, and centrifuged for 30 minutes at 4°C and 20,000 g. The supernatant was taken for Liquid Chromatography-Mass Spectrometry (LC-MS) analysis and ∼equal volume of supernatant was combined to make a QC sample and injected at the beginning and between every 6 samples to ensure instrument stability in day-to-day operation.

Metabolite separation was carried out at an acidic pH utilizing the Atlantis BEH Z-HILIC (2.1×150 mm, 2.5 M; part # 186009990; Waters Corporation) column and Thermo Scientific Vanquish Horizon UHPLC system. The mobile phase A (MPA) was 10 mM ammonium formate with 0.2% formic acid and mobile phase B (MPB) was acetonitrile with 0.1% formic acid. The parameters for the flow rate, injection volume, and column temperature were 30 °C, 5 L, and 0.2 mL/min, respectively. The chromatographic gradient was 0 minute: 90% B, 15 minutes: 20% B, 16 minutes: 20% B, 16.5 minutes: 90% B, 17 minutes: 90% B, and 23 minutes: 90% B. During the re-equilibration, the flow rate was increased to 0.4mL/minute for 4.7 minutes. Thermo Scientific’s Orbitrap IQ-X Tribrid mass spectrometer was used for MS detection, and it was equipped with an H-ESI probe that was working in positive polarity mode. The MS parameters were as follows: spray voltage: 3800 V, sheath gas: 80, auxiliary gas: 25, sweep gas: 1, ion transfer tube temperature: 300°C, vaporizer temperature: 300°C, automatic gain control (AGC) target: 25%, and a maximum injection time of 80 milliseconds (ms).

Thermo Scientific’s Xcalibur software was used to acquire the data in full-scan mode with a range of 70–1000 m/z at 60K resolution. The retention time and MS/MS fragmentation were compared to the reference standards in order to identify the metabolites. Data analysis was performed using Tracefinder 5.1 software (Thermo Scientific).

### Quantification and statistical analyses

All statistical analyses were performed using Prism 10 (GraphPad Software). Data were analyzed and indicated statistical comparisons for each statistical test are noted in each figure legend. Fold expansion curves, Fluo4/Fura Red curve and tumor growth curves were analyzed with repeated measures ANOVA. Data are presented as mean +/- standard error of the mean (S.E.M.) * p < 0.05; ** p < 0.01; **** p < 0.0001; ns, not significant.

## Data and code availability

The accession number for the RNA-seq data reported in this paper is GEO: GSE235214.

## Acknowledgements

The authors thank Dr. Creg J. Workman for the assistance with synapse imaging, Dr. Stacy G. Wendell and Dr. Steven J. Mullett for the help with lipidomics analysis and Mr. Leandro C. Hermida for the help with RNAseq analysis. AM was supported by the Brinson Foundation and the Cancer Research Foundation. We thank Dr. Lev Becker for sharing interstitial fluid samples from B16F10 murine tumors. We also thank Dr. Matthew Vander Heiden and Dr. Mark Sullivan for sharing interstitial fluid samples from BrafCA; PTENfl/fl; Tyr-CreER and MMTV-Cre; Brca1fl/fl; Trp53+/- tumor bearing mice.

---

**Supplementary Figure 1:**
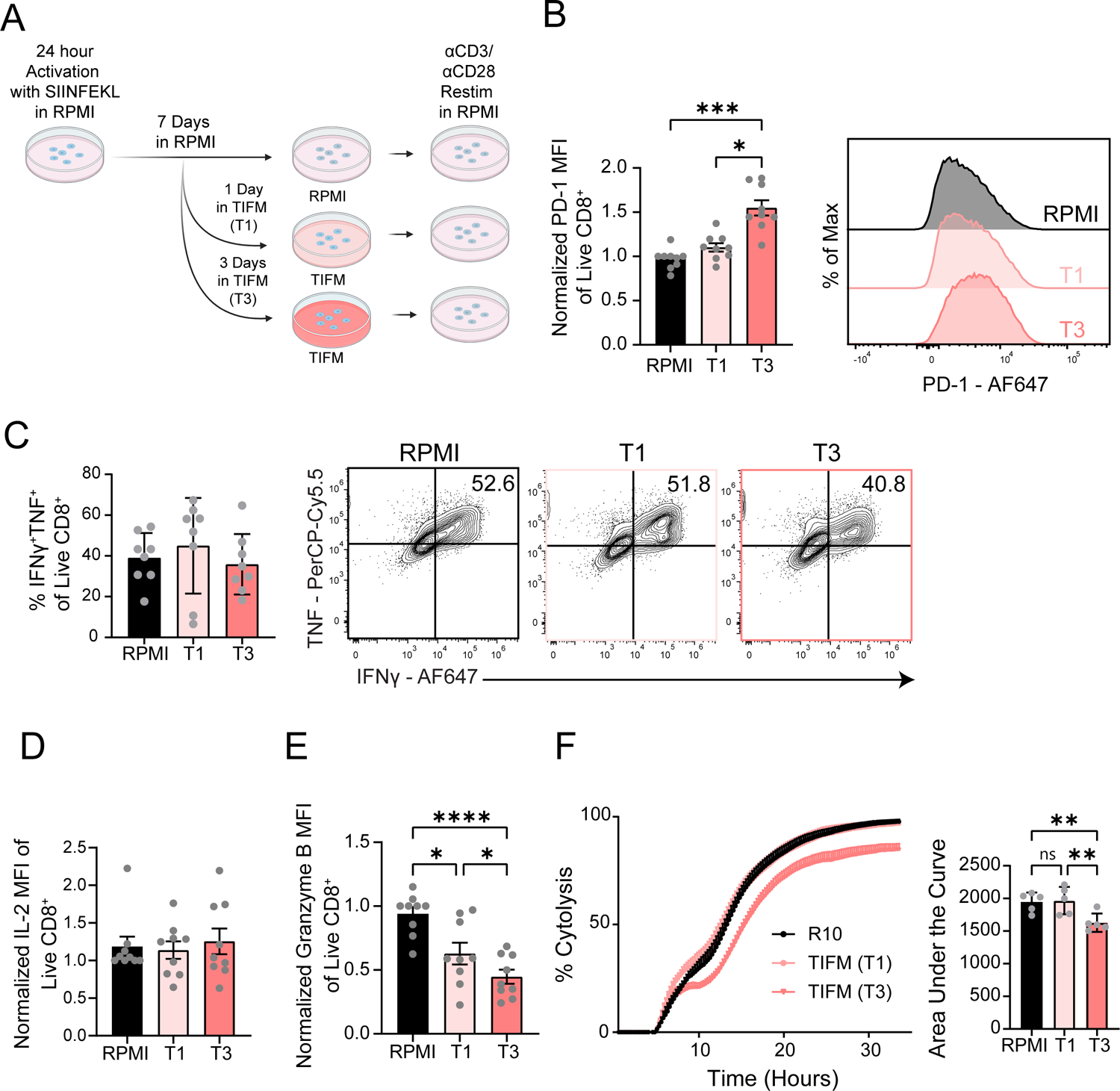
Shorter exposure to TIFM has effect on T cell inhibitory receptor function and Granzyme B expression **A)** Splenocytes from OT-I mice were activated with 250ng/mL SIINFEKL peptide for 24 hours in RPMI supplemented with 50U/mL IL-2. Cells were cultured for an additional 6 days in complete RPMI, or cells were cultured for 3 or 5 additional days in RPMI and then transferred into Tumor Interstitial Fluid Like Medium (TIFM) supplemented with IL-2 for 3 days or 1 day. For all panels in this figure, cells cultured in TIFM for the final 3 days of culture are labeled T3, and those cultured in TIFM for the final 1 day are T1. After 7 days, all cells were restimulated in RPMI with 3µg plate-bound αCD3 and 1µg soluble αCD28 antibody for 6 hours. **B)** Expression of PD-1 from Day 7 OT-I CD8^+^ T cells from indicated culture conditions and quantification of normalized PD-1 MFI (mean fluorescence intensity) ± SEM from four independent experiments is shown on the left (n=8-9/group). **C)** Day 7 OT-I CD8^+^ T cells were analyzed for IFNγ and TNF co-expression after 6 hours αCD3/αCD28 restimulation. Data quantified on the left show %IFNγ^+^TNF^+^ events among Live CD8^+^ T cells ± SEM from four independent experiments (n=8/group). **D-E)** Day 7 OT-I T cells were analyzed for expression of **D)** IL-2 or **E)** Granzyme B after 6 hours of αCD3/αCD28 restimulation. Data quantified on the left show normalized MFI ± SEM from four independent experiments (n=8-9/group). **F)** Day 7 OT-I CD8^+^ T cells were co-cultured with B16-OVA cells at a 1:1 effector:target ratio in an Agilent xCELLigence RTCA DP system to monitor target cell killing. XY plot on the left shows % Cytolysis ± SEM, and Area under the curve values are quantified on the right from two independent experiments (n=4-5/group). Statistical analysis for all indicated comparisons in **B-F)** was determined using a one-way analysis of variance (ANOVA) with Tukey’s multiple comparisons test. * = p<0.05, ** = p <0.01, *** = p<0.001, and **** = p<0.0001.

**Supplementary Figure 2:**
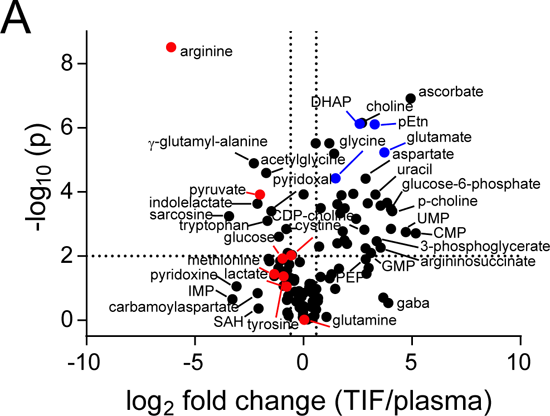
Differences in metabolite concentrations between TIF and plasma **A)** Fold change of the metabolite concentrations measured in the TIF relative to those in the plasma of mice bearing PDAC tumor, highlighted metabolites are marked by color to indicate metabolites that are depleted (red) and enriched (blue) in the tumor.

**Supplementary Figure 3:**
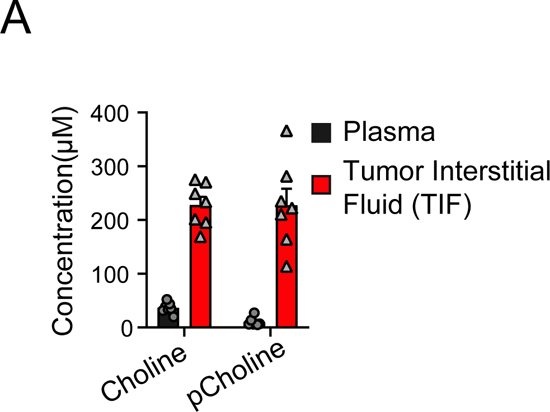
The TIF contains an altered metabolic milieu from that of the plasma **A)** Choline and phosphocholine levels in the plasma and the tumor interstitial fluid of mice bearing PDAC tumors. n=7/group.

**Supplementary Figure 4:**
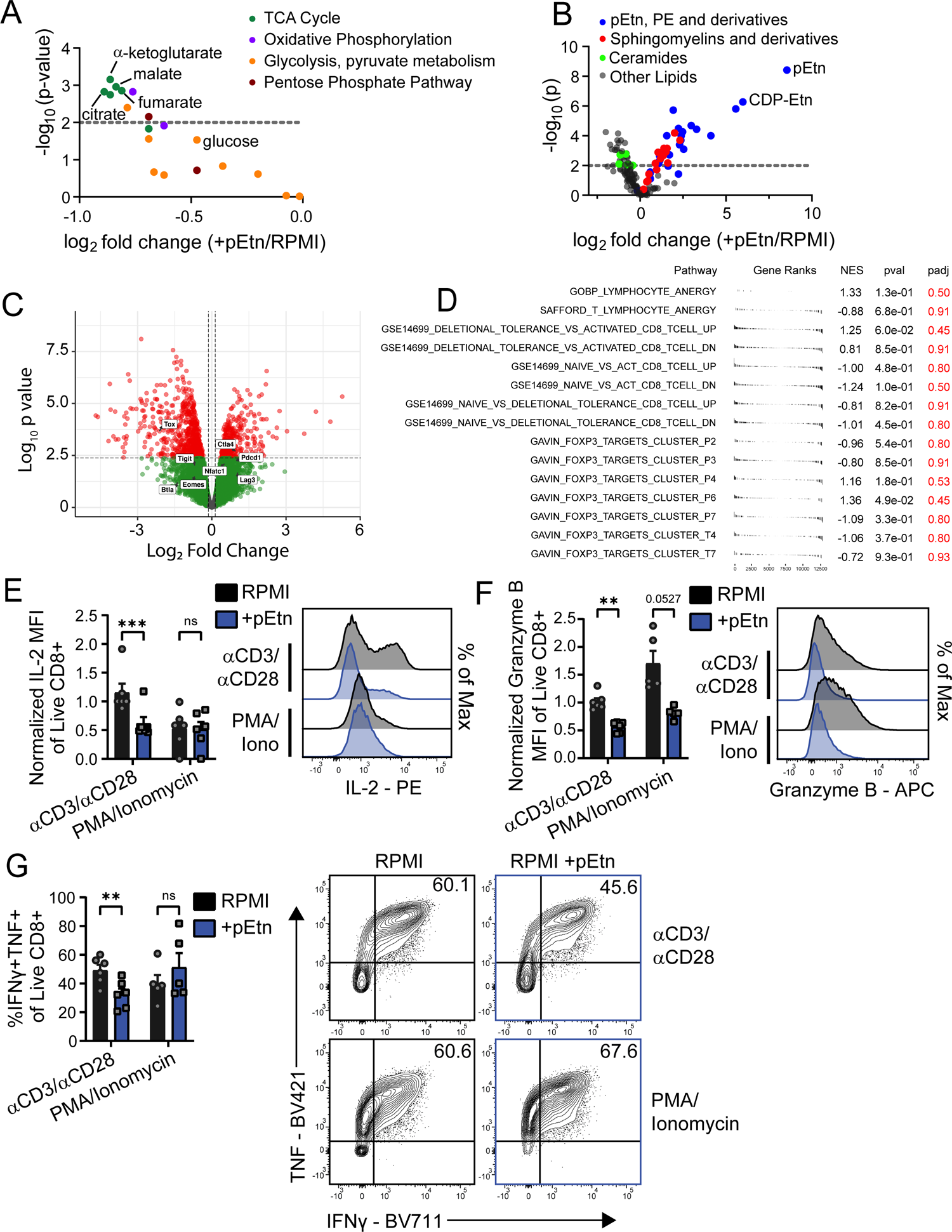
pEtn treated T cells have changes in metabolic and transcriptional pathways, and functional changes driven by pEtn can be rescued with PMA/Ionomycin. **A)** Quantification of metabolites in TCA cycle, oxidative phosphorylation, glycolysis, glutaminolysis and pentose phosphate pathway of SIINFEKL-activated OT-I T cells in RPMI+pEtn versus RPMI culture **B)** Quantification of lipid metabolites of SIINFEKL-activated OT-I T cells in RPMI+pEtn versus RPMI culture as shown in Figure 3A. **C)** Volcano plot of RNA-Sequencing of activated OT-I T cells cultured in RPMI or RPMI +pEtn. Highlighted genes represent exhaustion associated genes. **D)** Gene set enrichment assay of RNA-Seq data shown in **C)** of activated OT-I cells cultured in either RPMI or RPMI+pEtn. **(E-G)** OT-I T cells activated by SIINFEKL O/N were cultured in RPMI for 2 days and then in either RPMI or RPMI+pEtn for 5 days as described in Figure 3A). Then the cells were restimulated by either α-CD3 and α-CD28 or PMA/Ionomycin for 6 hours in RPMI. After restimulation, cells were analyzed for **E)** IFNγ and TNF co-expression **F)** Granzyme B expression, and **G)** IL-2 expression by flow cytometry. Normalized MFI ± SEM is quantified for Granzyme B and IL-2, while %IFNγ^+^TNF^+^ cells ± SEM are quantified on the left of each plot from three independent experiments (n=6/group). *p < 0.05, **p < 0.01, and ***p < 0.001 by paired t test. ns, non-significant.

**Supplementary Figure 5:**
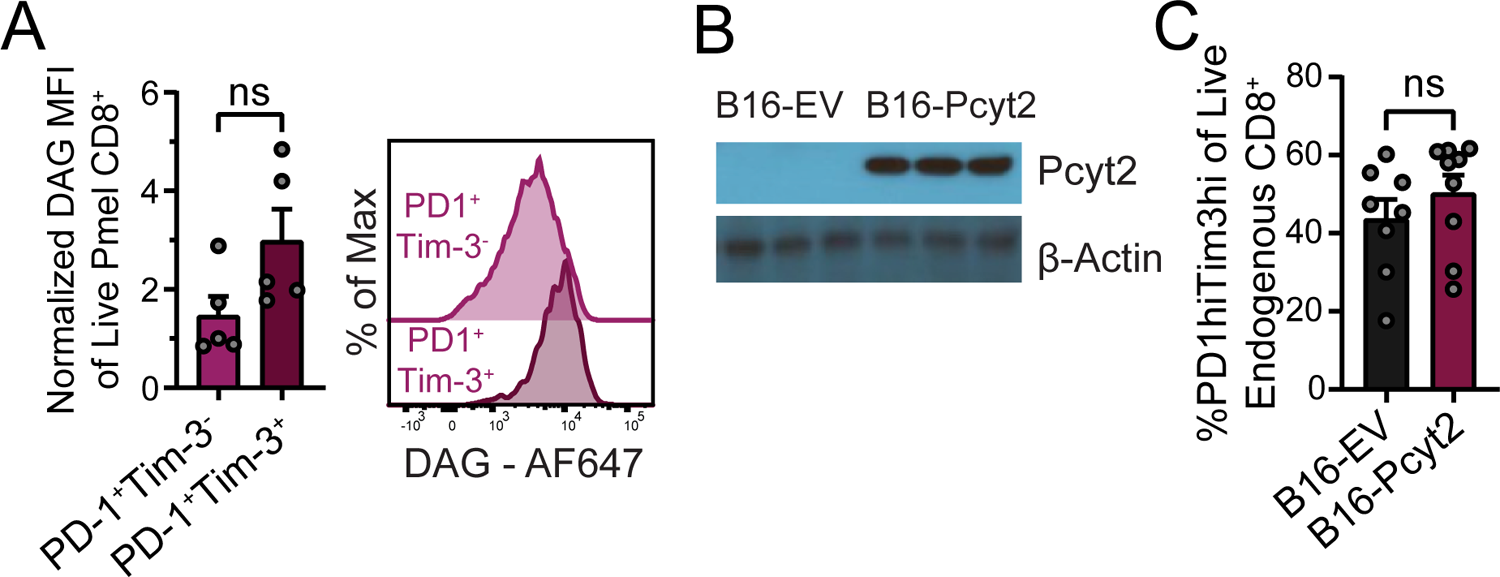
Pcyt2-overexpression B16 melanoma does not alter PD1^+^Tim-3^+^ frequency among the endogenous CD8^+^ T cells. **A)** DAG expression of the tumor infiltrating transferred Pmel T cells from the tumors in Fig 6A. Bar graph on the left shows normalized DAG MFI ± SEM from two independent experiments (n=5/group) **B)** Western blot for Pcty2 and Actin in B16 EV and B16-Pcyt2 melanoma cell lines. **C)** PD1 and Tim3 co-expression of endogenous CD8^+^ T cells from tumors in Fig 6I, Data quantified on the left show % PD1hiTim3hi from two independent experiments (n=11/group). Statistical significance for all indicated comparisons in this figure was calculated using an unpaired t-test.

**Table S1.**
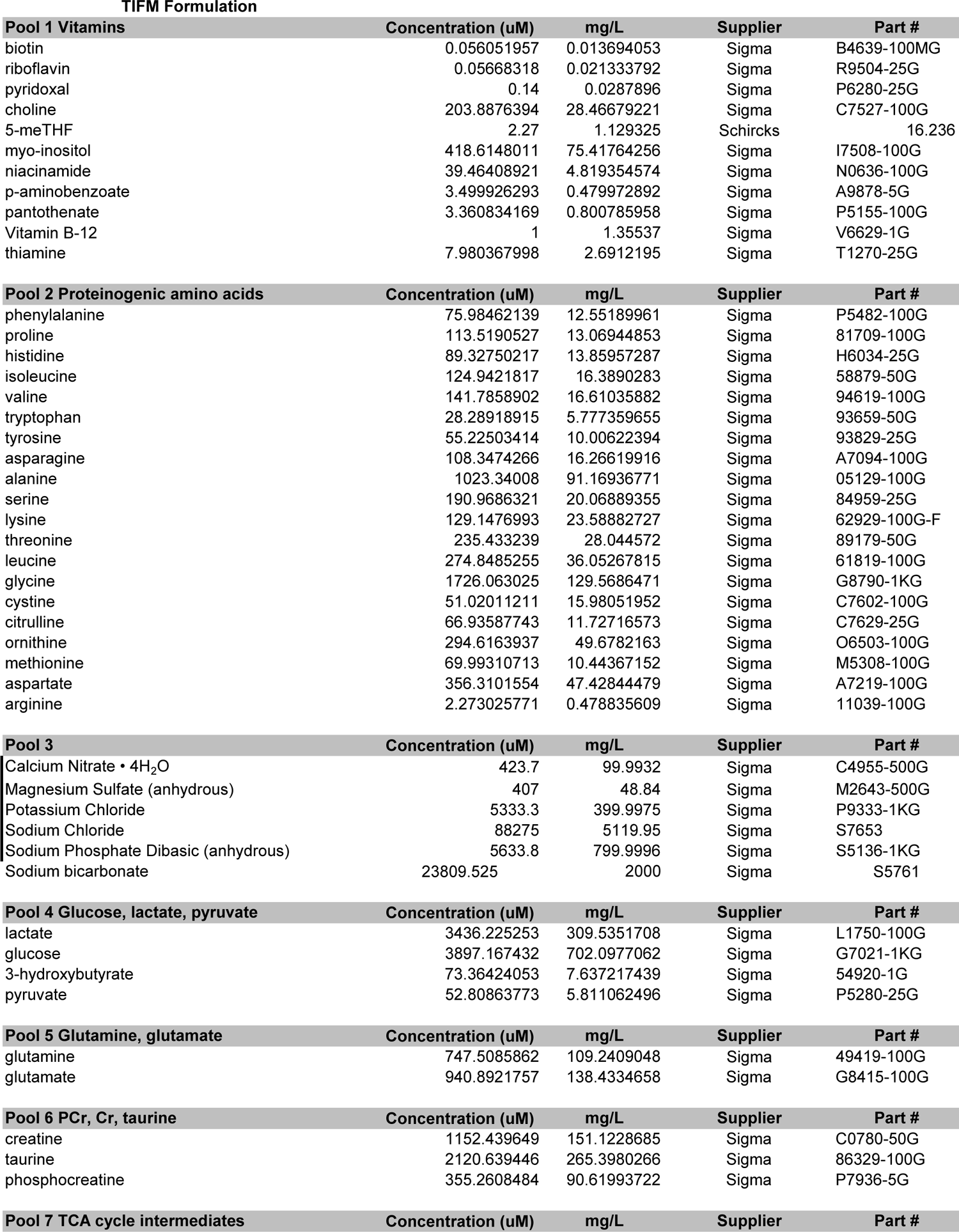
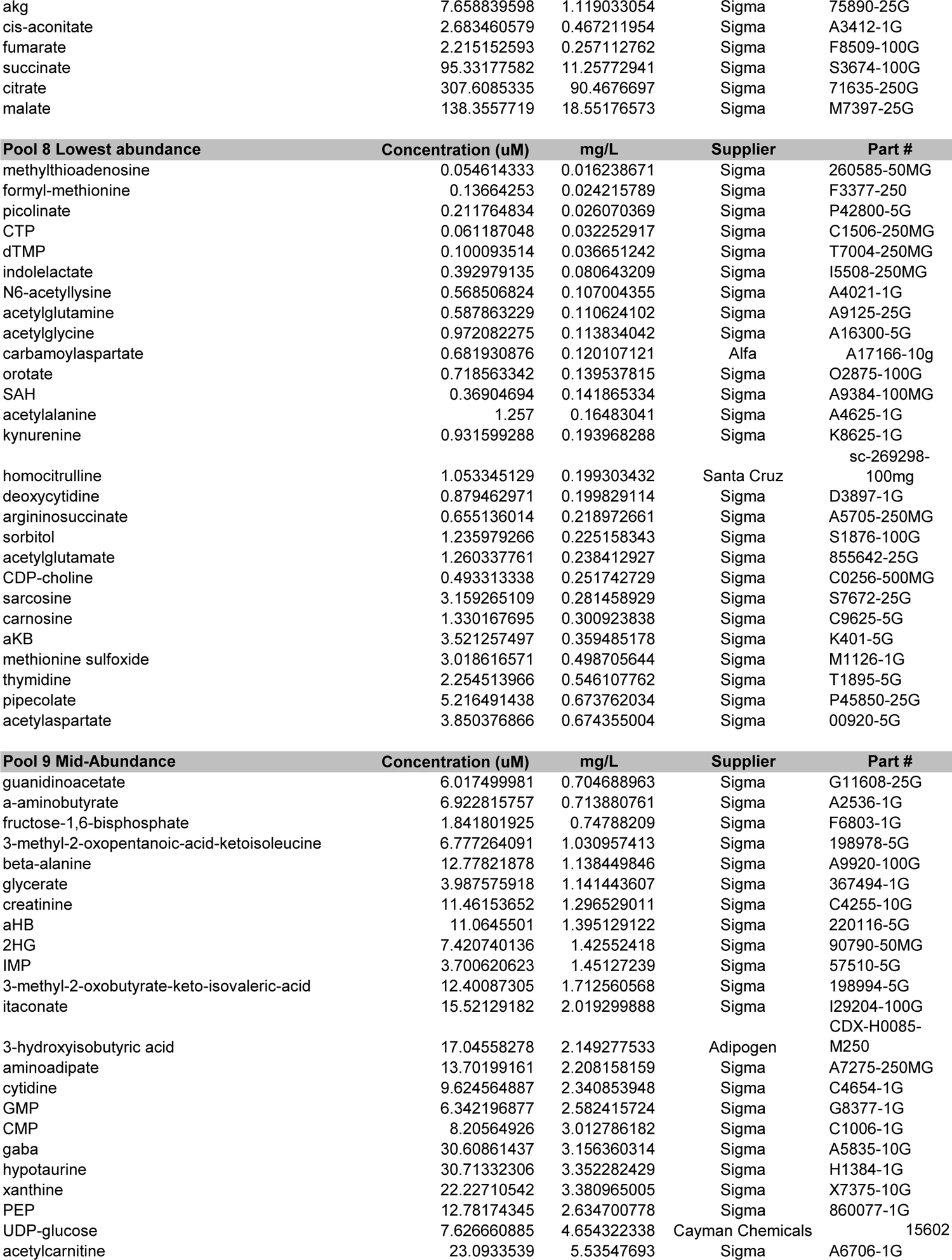
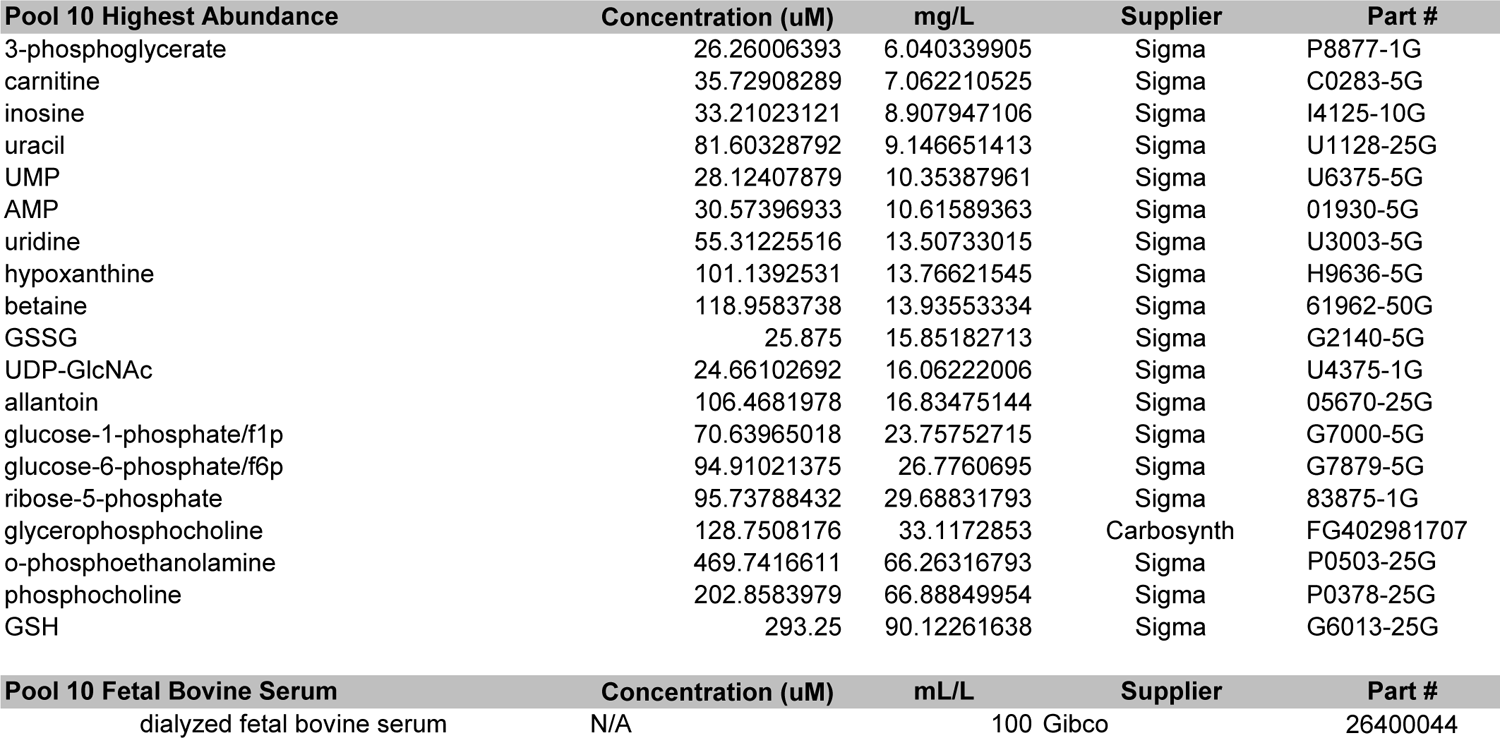

